# *Plasmodium* actin-like proteins are essential for DNA segregation during male gametogenesis and malaria transmission

**DOI:** 10.1101/2025.02.24.639803

**Authors:** Aastha Varshney, Eisha Pandey, Nirdosh, Satish Mishra

## Abstract

Protozoan parasites of the genus *Plasmodium* cause malaria and involve infection of multiple hosts and cell types during the life cycle. Producing sexually fit gametocytes is essential for transmitting the *Plasmodium* parasite into an anopheline mosquito vector. After the uptake of malaria parasites, male gametocytes undergo three rounds of DNA replication to produce eight nucleated flagellar gametes. Here, we report that the actin-like proteins Alp5a and Alp5b are involved in DNA segregation during male gametogenesis. The *Plasmodium*-specific paralogous genes Alp5a and Alp5b superimposed on human Arp2 and Arp3, localize to the nucleus, and interact with each other. Alp5a and Alp5b are dispensable for the development of *P. berghei* blood-stages individually but indispensable simultaneously. In agreement with genetic studies, the inhibitory activity of the Arp2/3 complex inhibitor in *Plasmodium* supports an essential role for this complex during the blood stage. Deletion of Alp5a or Alp5b did not affect actin nucleation, parasite growth, or gametocytemia in the blood. The knockout parasites successfully invaded the midgut and developed into oocysts, but their size was significantly reduced, and they failed to survive. Genetic crosses revealed defects in male gamete integrity. We found that the reduced oocyst development was due to impaired DNA segregation during male gametogenesis. This study provides molecular insight into the fundamental requirements of the Alps in *Plasmodium*, which are essential for malaria transmission.

## Introduction

Malaria parasites have a complex life cycle involving asexual and sexual replication in vertebrate and mosquito hosts. The sexual phase (gametogony) begins in the blood of the vertebrate host, whereas gametogenesis and meiosis require transmission to the mosquito host (1, 2). There are four periods of mitotic DNA synthesis and a single meiotic stage during the *Plasmodium* life cycle (1). In the blood stage, parasites proliferate via a process called schizogony. Mitotic cell division S during schizogony involves repeated rounds of nuclear division to form a multinucleated single cell (schizont), followed by one round of cellularization to form merozoites (3–5). The first cellular feature of schizogony is the formation of the hemispindle. The hemispindle is an array of microtubules that radiate into the nucleus from a single microtubule organizing center (MTOC) embedded in the nuclear membrane (6, 7). Once microtubules are formed, the spindle must assemble bipolarly to segregate the replicated chromosomes. The hemispindle retracts, and the MTOC duplicates to form a full, bidirectional mitotic spindle (8). Mitosis is regulated by cyclin-dependent kinases (CDKs), NIMA (Never In Mitosis)-related (NEK), and Aurora, Polo kinases (9–11).

The transmission of malaria parasites to their mosquito vectors relies upon the parasite switching from asexual reproduction to sexual reproduction (12). Male and female gametocytes produced during asexual schizogony are activated to begin gametogenesis to produce male and female gametes in the mosquito midgut (11). After activation, microgametocytes undergo three rounds of rapidly closed mitosis, increasing the DNA content from 1 N to 8 N. Concomitant spindle formation and chromosome segregation occur over 8–12 min without nuclear division, followed by karyokinesis and cytokinesis to bud eight microgametes (11, 13). Male gametogenesis is a complex process that is coordinated by well-established components such as mitogen-activated PK2 (MAPK2), serine-arginine PK1 (SRPK1) and calcium-dependent PK4 (CDPK4), as well as the metallo-dependent PP, PPM1 in *P. berghei* (11, 14–16). Other players include radial spoke protein 9 (RSP9) and Pb22, APC3 in *P. berghei* (5, 17, 18), p25α in *P. yoelii* (19) and the patatin-like phospholipase, PLP2 in *P. falciparum* (20). During male gametogenesis, rapid rounds of mitosis are driven by the formation of microtubular spindles to accurately segregate chromosomes. The kinetochore marker, NDC80, is located at distinct foci in the nucleus, and extends to form a bridge across the entire nuclear body during DNA segregation (21). The list of essential regulators of gametogenesis is reviewed in (11).

Actins are filament-forming, highly-conserved proteins essential for parasite development, host cell invasion, and male gametogenesis (22, 23). Malaria parasites express two actin isoforms, ubiquitous actin-1 and specialized actin-2 (24, 25). Actin-1 is an essential factor for *P. berghei* asexual blood stage development, and it cannot be deleted. Actin-2 is dispensable in the blood stage but is essential for male gametogenesis (26). In mutant parasites, male gametocyte DNA replicates normally, and axonemes assemble, but egress is severely blocked. This resulted in impaired exflagellation, and only a very small number of ookinetes were occasionally formed, but oocysts were not detected (26). Swapping of the actin-2 open reading frame (ORF) with actin-1 resulted in partial complementation of the defects in male gametogenesis. The complemented parasites produced ookinetes and were able to form oocysts, but these oocysts remained small, and their DNA was undetectable on day 8 post-blood meal. It does not develop into sporogonic oocysts (25). A report revealed that the F-actin isoform actin-2 forms within a few minutes after gametocyte activation and persists until the zygote transforms into an ookinete (27). Actin-2 is associated with the nucleus both in the male gametocyte and the zygote and readily forms long filaments necessary for male gametogenesis. Although actin-2 plays a specific, essential role in the maturation of male gametes, it polymerizes via the classical nucleation-elongation mechanism (23). An intriguing open question is the regulation of actin functions in the absence of actin regulatory proteins in *Plasmodium*. Isodesmic polymerization of consecutive monomers has been suggested for *T. gondii* actin, which differs from the classic nucleation of actins (28). However, a study reported a nucleation-elongation mechanism of actin in *Plasmodium* rather than an isodesmic mechanism (28). The actin nucleation factor Arp2/3 complex is characterized in other organisms. Malaria parasites have a minimal set of proteins that potentially regulate microfilament dynamics and lack canonical actin-nucleating factors such as the Arp2/3 complexes (29). In *Plasmodium*, the formin family of proteins is thought to represent the only actin nucleators (30). The limited sequence conservation of *Plasmodium* proteins is one of the reasons for knowledge gaps (31). In fact, many proteins are currently considered absent or referred to as uncharacterized proteins in *Plasmodium*, such as the actin nucleator actin-related protein 2/3 (Arp2/3) complex (32–35).

The Arp2/3 complex nucleates actin filaments and comprises seven proteins with five unique polypeptides, called ARPC1-5, supporting a dimer of two Arps (actin-related proteins Arp2 and Arp3) (32, 33, 36). The complex is well known to be conserved throughout the eukaryotic kingdom. However, its function has been validated in a few model organisms (37). The Arp2/3 complex plays a role in actin polymerization, coordinating the nucleation of spindle actin during mitosis and meiosis and accurate chromosome segregation (38–41). The Arp2/3 complex itself has been conserved throughout evolution (42, 43), and it was assumed that the complex has been lost in apicomplexan parasites except ARPC1/ARC40 in malaria parasites (34, 35). Here, we found that Alp5a and Alp5b are related to human Arp2 and Arp3, and the STRING database revealed a noncanonical Arp2/3 complex in *Plasmodium*. We analyzed the expression and function of *P. berghei* Alp5a and Alp5b. The Alp5 KO parasites could form ookinetes, but the resulting oocysts presented a delayed death phenotype, leading to a block in further malaria transmission. Genetic crosses revealed a defect in the male gamete in Alp5 KO parasites. The defect was due to impaired DNA segregation during male gametogenesis.

## Results

### *Plasmodium* Alp5a and Alp5b are structurally similar to human Arp2 and Arp3, localize to the nucleus, and interact with each other

We started our study by in silico analysis of the paralogous genes Alp5a and Alp5b. NCBI BLASTP analysis of *P. berghei* Alp5s revealed that Alp5a and Alp5b are closely associated with *P. yoelii* and *P. falciparum,* respectively (Figure S1). The protein-protein interactions of both Alp5a and Alp5b were confirmed via the STRING database. An interactome of 11 proteins revealed that all the protein interactions were the same for Alp5a and Alp5b, except for PBANKA_1434600 (formin2, putative protein) in Alp5a (Figure S2). All the interacting proteins are conventional or actin-like, dynactin subunits, formins, or subunits of the Arp2/3 complex. This indicated the presence of a noncanonical Arp2/3 complex (Figure S2). The ClusPro server predicted high-affinity interactions of Alp5 with actin (Figure S3). This prediction suggests that Alp5a and Alp5b interact with conventional actin and play roles in actin dynamics and the polymerization of actin filaments. To assess the interaction of Alp5 with actin, we generated a 3D model of actin isoforms (actin 1 and actin 2) (Figure S3A and D). Cluspro protein-protein docking studies indicated that Alp5a has a stronger affinity for both actin isoforms than Alp5b (Figure S3B, C, E and F). The predicted protein structures of Alp5a and Alp5b were generated via Alphafold2 and assessed via UCLAsaves, and the best structures were selected (Figure 1A and B). Alp5a and Alp5b were superposed with the human Arp3 and Arp2 structures via PyMOL. We observed the superposition of Alp5a and Alp5b with human Arp3 and Arp2, with RMSD values of 2.960 and 1.328, respectively (Figure 1C and D). Next, Alp5a and Alp5b were checked for interactions via the ClusPro server and visualized via PyMOL. The interaction scores suggested their possible interactions (Figure 1E).

**Figure 1.**
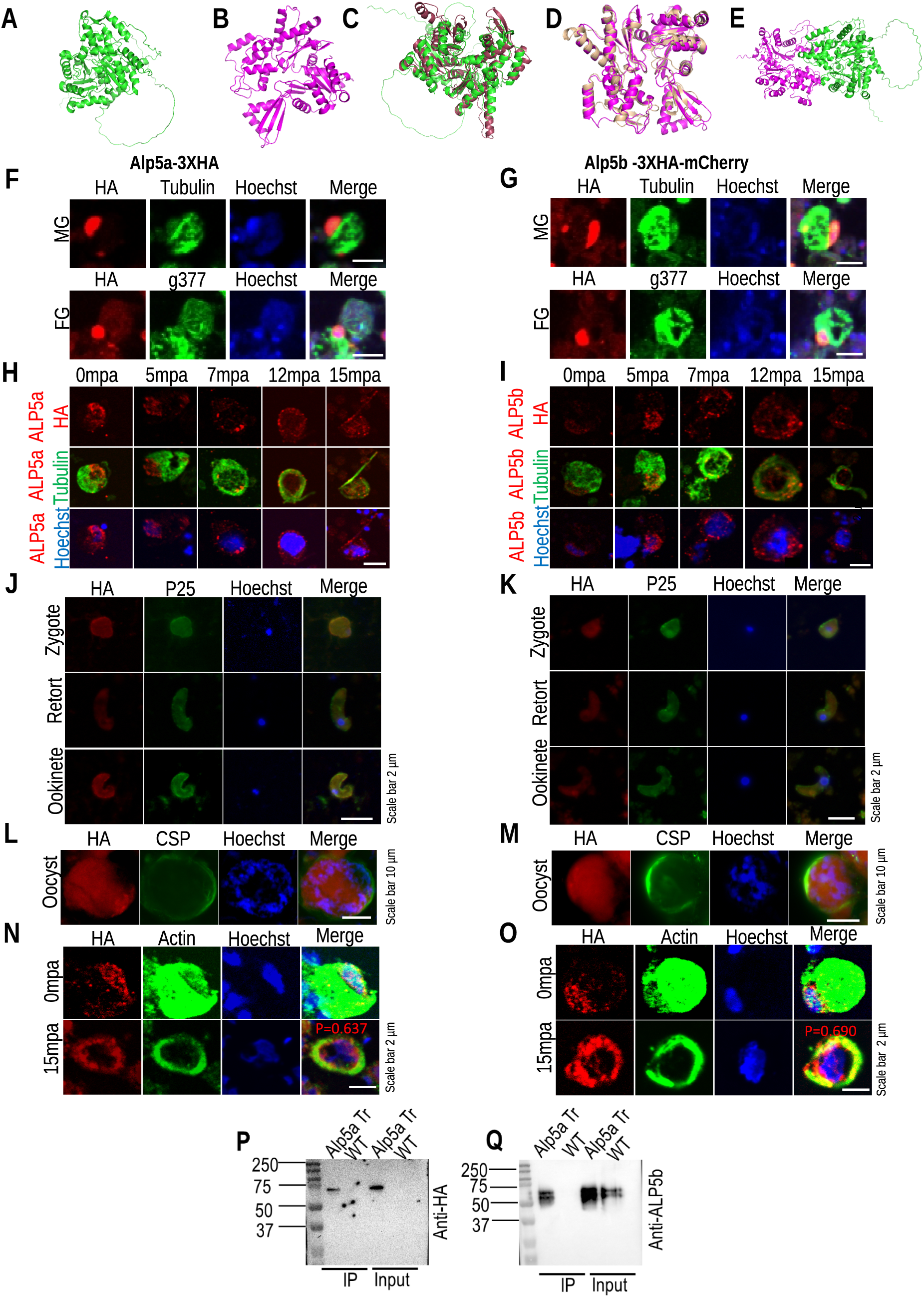
Expression and localization of Alp5s and their interaction. **(A)** The cartoon structure of *Plasmodium berghei* Alp5a was visualized via PyMOL. The diagram simplifies the protein’s secondary structure, highlighting its overall fold and key structural features. **(B)** Cartoon structure of Alp5b. **(C)** The structural superposition of *Plasmodium berghei* Alp5a with human Arp3 shows an RMSD (root mean square deviation) value of 2.960 Å. This RMSD value quantifies the structural alignment between the two proteins, illustrating the degree of similarity and divergence in their conformations. **(D)** The structure superposition compares *Plasmodium* Alp5b with human Arp2 with an RMSD value of 1.328. This comparison highlights the similarities and differences in their structural conformations, providing insights into potential functional parallels and divergences between the *Plasmodium* protein and its human counterpart. **(E)** The interaction between *Plasmodium berghei* Alp5a and 5b, as predicted by ClusPro. The docking model illustrates the predicted binding interface and interaction sites between the two proteins, highlighting their potential functional relationships. **(F and G)** Representative immunofluorescence images of male and female gametocytes immunostained with anti-HA, anti-tubulin and anti-g377 antibodies. The Alp5a-3XHA and Alp5b-3XHA-mCherry parasites show their localization. Nuclei were stained with Hoechst. DIC: differential interference contrast. **(H and I)** Immunofluorescence analyses of male gametes after gametocyte activation. The cells were fixed and immunostained with anti-HA and anti-tubulin antibodies. Alp5a and Alp5b localized from the nucleus to axonemes. **(J and K)** Immunofluorescence analyses of Alp5a-3XHA and Alp5b-3XHA-mCherry revealed their localization at the zygote, retort, and ookinete stages. Parasites were identified by immunostaining with an anti-P25 antibody. **(L and M)** Expression of Alp5a-3XHA and Alp5b-3XHA-mCherry parasites in oocysts. The midgut oocysts were labeled with anti-HA and anti-CSP antibodies. **(N and O)** Male gametocytes of Alp5 transgenic parasites were immunostained with anti-HA and anti-actin antibodies at 0- and 15-min post-activation. At 0 mpa, Alp5 was localized mainly in the nucleus, whereas actin was distributed in cytosol and nucleus. Post-activation at 15 mpa, Alp5, and actin are redistributed around the nucleus and colocalize with each other. **(P and Q)** Interaction of Alp5s in the parasite. The complex was immunoprecipitated from Alp5a-3XHA transgenic parasites via anti-HA magnetic beads. The Alp5a and Alp5b bands were detected by immunoblotting with anti-HA and anti-Alp5b antibodies.

Next, to check the expression, localization, and interactions of Alp5, we endogenously tagged Alp5a and Alp5b with 3XHA and 3XHA-mCherry, respectively (Figure S4A, B, D, and E). Western blot analysis with an anti-HA antibody confirmed the expression of correctly sized Alp5 fusion proteins in the gametocytes of the transgenic parasites (Figure S4C and F). IFA revealed the expression of Alp5a and Alp5b in male and female gametocytes but not in the ring or trophozoite stages. Male and female gametocytes were immunostained with anti-tubulin and anti-g377 antibodies to confirm gamete-specific expression. Both Alp5a and Alp5b were expressed predominantly in the nuclei of gametocytes (Figure 1F and G). The signals of Alp5a and Alp5b localize from the nucleus to the axoneme (Figure 1H and I). Diffuse signals were also observed in the parasite cytoplasm. The expression of Alp5a and Alp5b was also detected in zygotes, ookinetes, and oocysts (Figure 1J-M). The expression of Alp5a and Alp5b was not detected in sporozoites or exo-erythrocytic forms (Figure S5A and B). To assess the colocalization of Alp5 and actin, nonactivated and activated gametocytes were immunostained with anti-HA and anti-actin antibodies. The colocalization of Alp5 and actin in activated gametocytes indicated their possible interaction (Figure 1N and O). To determine the endogenous interaction between Alp5a and Alp5b, the complex was immunoprecipitated from gametocytes of Alp5a-3XHA transgenic parasites using anti-HA magnetic beads. The blot was probed with an anti-HA antibody that detected the Alp5a band (Figure 1P). The blot was reprobed with an anti-Alp5b antibody, which detected the Alp5b band, indicating the interactions of Alp5a with Alp5b (Figure 1Q). These results suggest that Alp5 is predominantly expressed in the sexual stages and that the Alp5a and Alp5b subunits interact with each other.

### ALP5a and Alp5b are individually dispensable in the blood, but the Arp2/3 complex is essential for parasite survival

Alp5a and Alp5b are paralogous genes; hence, we studied their roles together. To understand the role of Alp5a and Alp5b in the parasite life cycle, we disrupted the gene in *P. berghei* via double crossover homologous recombination (Figure S6A, B, E and F). To restore gene function in the KO parasites, a complemented line was generated by reintroducing the Alp expression cassette (Figure S6C, D, G, H and I). We attempted to disrupt both genes simultaneously to check for redundancy in the functions of Alp5a and Alp5b. We applied two independent strategies for the double disruption of genes. First, we transfected an Alp5b-targeting construct into Alp5a KO schizonts. (Figure S7A). Twenty-four hours post-transfection, we observed a few parasites with red and green fluorescence, which were lost after WR drug selection. Next, we transfected *P. berghei* WT schizonts with Alp5a and Alp5b targeting constructs (Figure S7B). Parasites were selected using pyrimethamine drugs. We observed individual KO parasites, but no double-deletion events were observed (Figure 2A). We treated parasites with different inhibitors to further confirm the essentiality of actin and the Arp2/3 complex in *Plasmodium* blood stages. We found that an Arp2/3 complex inhibitor (CK-666), an actin polymerization inhibitor (Cytochalasin D), and an actin polymerization inducer (Jasplakinolide) inhibited the growth and development of *P. falciparum* and *P. berghei* parasites (Figure 2B, C, S8A and B).

**Figure 2.**
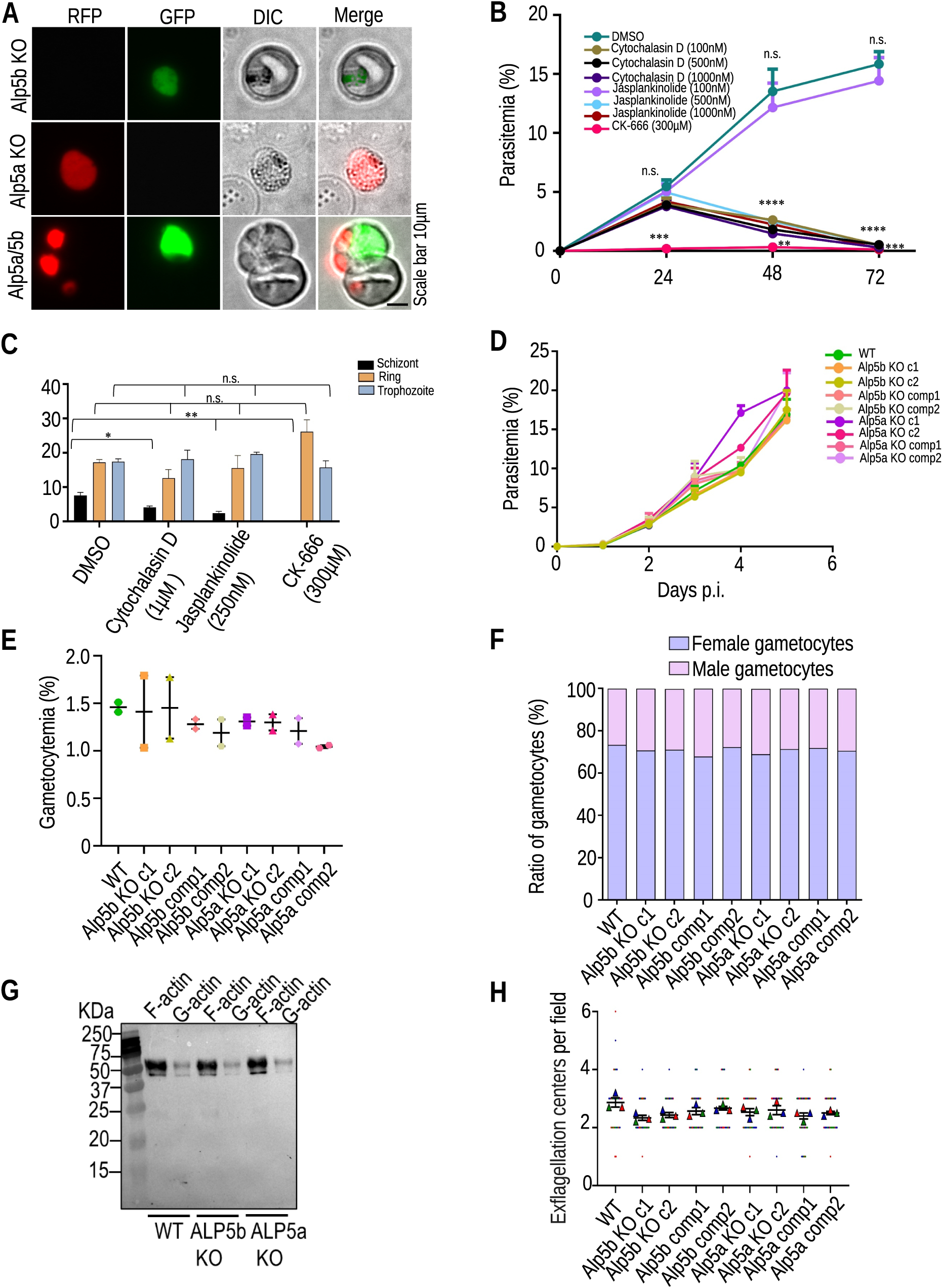
Alp5a and Alp5b are not individually required for asexual blood stage propagation, but simultaneously, they are indispensable. **(A)** mCherry- or GFP-expressing individual Alp5a- or Alp5b KO parasites survived after drug selection. Even independent KO parasites survived within the same RBC, but parasites with double deletions were not observed. **(B)** Effect of inhibitors on the growth of *P. falciparum.* Parasitemia at 24, 48 and 72 h after treatment with the notated doses of cytochalasin D, jasplankinolide and CK-666. We treated synchronized *P. falciparum* cultures in triplicate. There was no difference in growth at 24 h in the presence of DMSO vs cytochalasin D or jasplankinolide (P=0.2631; one-way ANOVA). Inhibition was observed in the CK-666-treated cultures, DMSO vs CK-666, ***P = 0.0008; unpaired Student’s t test. A reduction in growth was observed at 48 h, DMSO vs cytochalasin D or jasplankinolide (500 nm-1 µM) ****P<0.0001; one-way ANOVA test, DMSO vs CK-666, **P =0.0021; unpaired Student’s t test. The growth further decreased at 72 h. DMSO vs cytochalasin D or jasplankinolide (500 nm-1 µM), ****P<0.0001 and DMSO vs CK-666 ***P =0.0001; unpaired Student’s t test. Compared with DMSO, 100 nM jasplankinolide did not affect parasite growth at 48 h (P= 0.6447) or 72 h (P= 0.5590). The data are presented as the mean ± SEM and were analyzed via unpaired Student’s t tests; n=3 biological replicates. **(C)** *P. berghei* blood cultures were treated with the inhibitors cytochalasin D (1 µM), jasplankinolide (250 nM) and CK-666 (300 µM). Counting of ring, trophozoite, and schizont stages in *P. berghei* cultures after treatment. There was no effect on the ring (cytochalasin D, P= 0.1605; jasplankinolide, P= 0.6896; and CK-666, 0.0693) or trophozoite stage (cytochalasin D, P= 0.8053; jasplankinolide, P= 0.1031; and CK-666 (P=0.4832. The number of schizonts decreased significantly in all the treated groups (cytochalasin D, *P= 0.0257; jasplankinolide, **P= 0.0082; and CK-666, **P=0.0012). **(D)** Asexual blood-stage propagation of control and KO parasites. The data are presented as the mean ± SEM, with no significant differences (P=0.9998; Brown-Forsythe ANOVA)**. (E)** Estimation of gametocytemia. The data are presented as the mean ± SEM with no significant differences (P=0.7776; Brown-Forsythe ANOVA test). **(F)** The male and female gametocyte ratios were comparable in WT and KO parasites. The data are presented as the mean ± SEM, with no significant differences (P=0.9864; one-way ANOVA). Data from two independent experiments are presented in D, E, and F. **(G)** Western blot analysis of F-actin and G-actin in blood-stage parasites. Parasites were homogenized in F-actin stabilization buffer, followed by centrifugation to separate F- actin from the G-actin pool. Immunoblotting was performed with an anti-actin antibody to detect actin. Western blot quantitation of F-actin and G-actin in blood-stage parasites and data are presented as the F/G actin ratio. There was no difference between WT and Alp5 KO parasites (P=0.5903; one-way ANOVA). **(H)** Comparison of exflagellation centers in WT and Alp5 KO parasites revealed no difference (P= 0.1274, Brown Forsythe and Welch ANOVA). Counts were checked from 10 random fields in three independent experiments. The data are presented as the mean ± SEM.

For phenotypic analysis of Alp5a and Alp5b KO parasites, we assessed asexual blood stage propagation. Both KO parasites propagated normally and produced gametocytes at a similar rate (Figure 2D and E). Next, we examined the proportions of male and female gametocytes and found that they were comparable in WT and KO parasites (Figure 2F). We also analyzed the F-actin/G-actin ratio in the KO parasites, which was comparable to that in the WT parasites (Figure 2G). Next, we analyzed microgamete formation. No significant differences were observed in the number of exflagellation centers of KO and WT parasites (Figure 2H). These results indicate that the Alps are not individually required for actin nucleation or blood-stage development but are essential as a complex for parasite survival.

### Alp5s are essential for male gamete fertility, oocyst development, and malaria transmission

To analyze Alp5 KO parasite development in mosquitoes, we transmitted parasites by feeding them to infected mice. We observed oocyst development on day 14 post-bloodmeal and found a significant reduction in oocyst size in both types of KO parasites (Figure 3A). Next, we analyzed the size and number of oocysts from days 8 to 18 post-feeding. The size and number of oocysts were significantly reduced in KO parasites. Interestingly, the number of oocysts decreased over time, and only a few remained on day 18 (Figure 3B and C). Oocyst size did not increase, and sporogony was abolished in the KO parasites (Figure 3D). We crushed the midgut and salivary glands to estimate the sporozoite numbers. The sporozoites were completely absent in the Alp5 KO parasites (Figure 3E, F and G). All the KO-infected mosquitoes failed to transmit malaria to the mice in the biteback experiment (Figure 3H and Table 1). The complemented parasite lines successfully completed all the developmental stages, confirming that the observed phenotype was caused by the deletion of the Alp5 gene (Figure 3). These results demonstrate that Alp5 is required for oocyst development and sporogony and that deleting Alp5s blocks malaria transmission.

**Figure 3.**
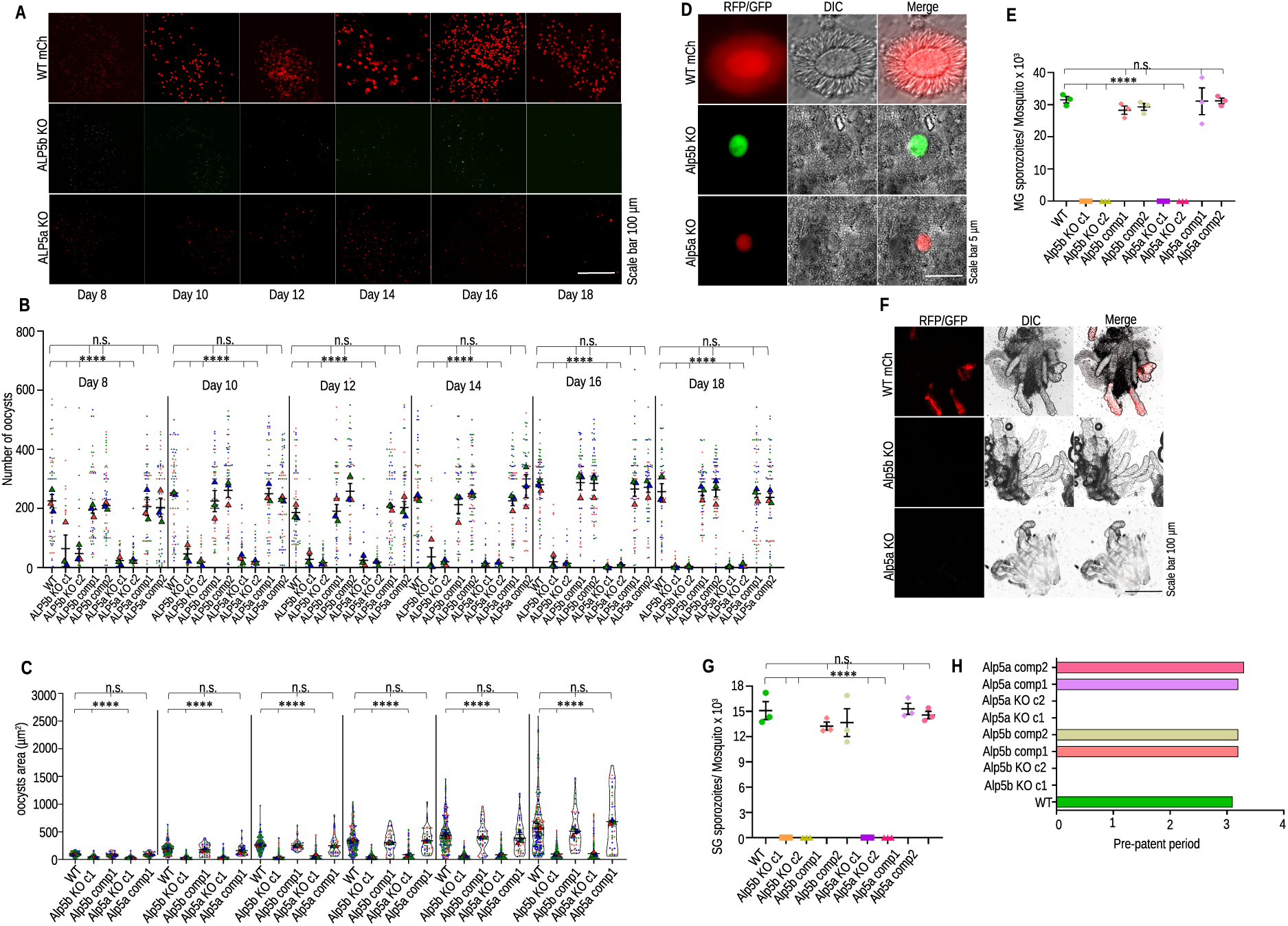
Alp5s are essential for oocyst development and malaria transmission. Mosquito midgut showing oocysts on the indicated days. **(B)** Quantification of oocyst numbers. There was a significant difference in oocyst number between WT and KO parasites on days 8-18 (****P<0.0001). In contrast, no differences were detected between the complement lines and WT on days 8 (P= 0.8721), 10 (P= 0.5445), 12 (P=0.0662), 14 (P=0.1075), 16 (P=0.7252) or 18 (P=0.8941). Sixty midguts from all the groups were dissected each day. **(C)** Determination of the oocyst area. The data from independent clones were pooled, and a significant difference was detected between the WT and KO lines on days 8-18 (****P<0.0001), whereas no differences were detected between the complemented lines and the WT line on days 8 (P= 0.1058), 10 (P= 0.0890), (P=0.1553), 14 (P=0.8983), 16 (P=0.5481) or 18 (P=0.5198). The Kruskal‒Wallis test was used to determine the significance. Two hundred oocyst areas in the WT, Alp5a KO, and Alp5b KO lines and 40 oocyst areas in the complement lines were compared on the indicated days. Kruskal‒Wallis tests were performed, and the data represent three independent experiments. The triangle represents the mean individual count in each experiment, and each dot represents an individual count. **(D)** Oocysts showing sporogony in WT. No sporogony was observed in Alp5a or Alp5b KO oocysts. **(E)** Midgut sporozoite count. The data revealed a significant difference between the WT and knockout lines (****P<0.0001). No difference was observed between the WT and complemented lines (P=0.7737). The total number of mosquitoes dissected in each group was 95 (WT GFP), 235 (Alp5b c1), 235 (Alp5b c2), 95 (Alp5b comp1), 70 (Alp5b comp2), 260 (Alp5a c1), 245 (Alp5a c2), 80 (Alp5a comp1), and 85 (Alp5a comp2). **(F)** Salivary gland sporozoites were observed in the WT but not in the KO-infected glands. **(G)** Salivary gland sporozoite count. The total number of mosquitoes dissected in each group was 85 (WT GFP), 118 (Alp5b c1), 170 (Alp5b c2), 70 (Alp5b comp1), 100 (Alp5b c2), 135 (Alp5a c1), 235 (Alp5a c2), 70 (Alp5a comp1), and 70 (Alp5a comp2). The data revealed a significant difference between the WT and knockout lines (****P<0.0001). No difference was detected between the WT and complemented lines (P=0.5317). **(H)** Transmission of sporozoites in mice by mosquito bites. Infection was not observed in the mice inoculated with knockout-infected mosquitoes. One-way ANOVA was performed to determine the significance. The data were pooled from three independent experiments.

**Table 1.** In vivo Infectivity of Alp5 KO parasites in C57BL/6 mice. Sporozoites were inoculated into C57BL/6 mice via mosquito bites or intravenous injection. The presence of parasites in the blood was confirmed via a Giemsa-stained blood smear.

| Experiment | Parasites | Number of mosquitoes/mouse | Mice positive/Mice infected | Pre-patent period (days) |
| --- | --- | --- | --- | --- |
| **1** | WT-mCherry | 50 | 5/5 | 3.2 |
|  | Alp5b KO c1 | 50 | 0/5 | NA |
|  | Alp5b KO c2 | 50 | 0/5 | NA |
|  | Alp5b comp1 | 50 | 5/5 | 3.2 |
|  | Alp5b comp2 | 50 | 5/5 | 3.4 |
|  | Alp5a KO c1 | 50 | 0/5 | NA |
|  | Alp5a KO c2 | 50 | 0/5 | NA |
|  | Alp5a comp1 | 50 | 5/5 | 3.4 |
|  | Alp5a comp2 | 50 | 5/5 | 3.2 |
| **2** | WT | 50 | 5/5 | 3 |
|  | Alp5b KO c1 | 50 | 0/5 | NA |
|  | Alp5b KO c2 | 50 | 0/5 | NA |
|  | Alp5b comp1 | 50 | 5/5 | 3.2 |
|  | Alp5b comp2 | 50 | 5/5 | 3 |
|  | Alp5a KO c1 | 50 | 0/5 | NA |
|  | Alp5a KO c2 | 50 | 0/5 | NA |
|  | Alp5a comp1 | 50 | 5/5 | 3 |
|  | Alp5a comp2 | 50 | 5/5 | 3.4 |
| Experiment | Parasites | Number of sporozoites injected | Mice positive/Mice injected | Prepatent period (days) |
| **1** | WT | 5,000 | 5/5 | 3.2 |
|  | WT-mCh×Alp5b KO | 5,000 | 5/5 | 3.4 |
|  | WT-GFP×Alp5a KO | 5,000 | 5/5 | 3.6 |
|  | Pb47×Alp5a KO | 5,000 | 5/5 | 3.6 |
|  | Pb47×Alp5a KO | 5,000 | 5/5 | 3.6 |

After the essential roles of Alp5a and Alp5b in oocyst development and malaria transmission were determined, ookinete formation was analyzed. We found no difference in ookinete number or morphology between WT and KO parasites (Figure S9A and B). A genetic cross was performed between the KO and WT parasites to analyze sex-specific defects in the KO parasites. On day 15, post-feeding, we dissected the midgut, and the development of oocysts was observed. Genetic crosses with WT parasites restored the KO parasite’s phenotype (Figure 4A), indicating a defect at the gamete stage. *Plasmodium* inherits many defects during mosquito stage development from one sex (11). To investigate whether the functions of Alp5a and Alp5b are sex specific, we generated male and female gamete defective lines by knocking out the Pb48/45 (44) and Pb47 genes (45), respectively (Figure S10). Next, we crossed KO parasites with gamete defective lines (Pb47×Alp5b KO, Pb48/45×Alp5b KO, Pb47×Alp5a, Pb48/45×Alp5a). The Alp5 KO × Pb47 cross showed robust oocyst development, indicating that fertile female gametes were present in the Alp5 KO parasites (Figure 4B and S11). In contrast, the Alp5 KO×Pb48/45 cross showed no change in oocyst development, indicating defects in male gamete function in Alp5 KO parasites (Figure 4B and S11). The oocysts were counted, areas were measured, and sporogony was observed after the genetic cross-experiments. The sporogony of the restored oocysts was normal (Figure 4C and D). On day 19, post-feeding, we dissected the mosquitoes and detected a normal sporozoite load in the salivary glands of all the groups except the Alp5-KO lines, which were crossed with a male gamete-defective line (Figure S12A and B). The obtained salivary gland sporozoites were checked for infectivity in vivo and in vitro. The sporozoites were injected into C57BL/6 mice, and the pre-patent period was observed beginning on day three post-infection (Table 1). The groups with restored salivary gland sporozoite loads infected hepatocytes and transitioned to the blood stage normally (Figure S13). Together, these results demonstrated that Alp5s play male-specific roles in gamete fertility.

**Figure 4.**
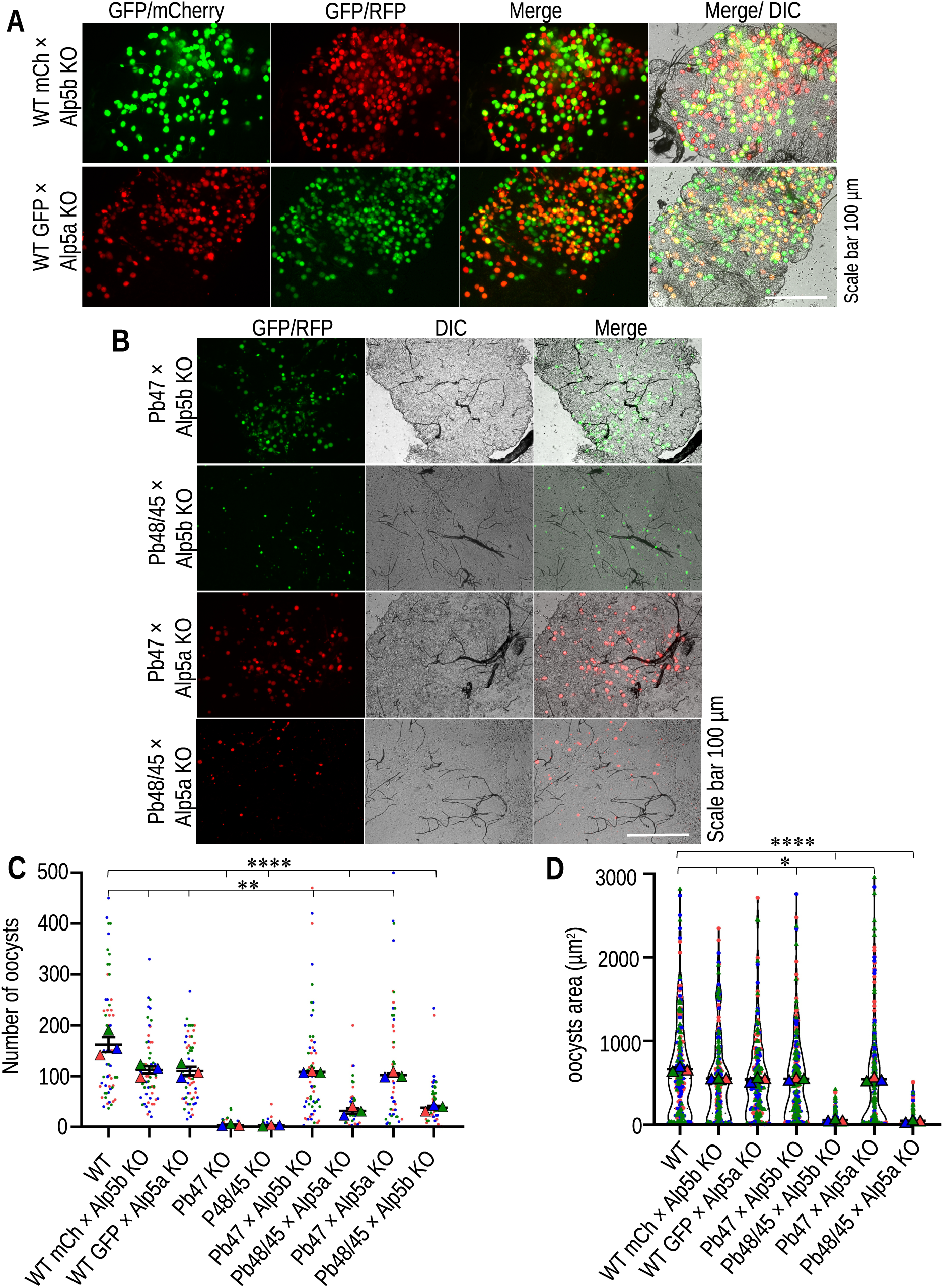
Genetic cross revealed a defect in the male gamete in Alp5 KO parasites. **(A)** Observation of oocysts on day 15 post-feeding after genetic crosses with WT-mCherry×Alp5b KO or WT-GFP×Alp5a KO parasites. **(B)** Genetic crosses of Alp5 KO parasites with male or female gamete-defective lines. The phenotype was restored in the KO lines crossed with the female gamete-defective lines. Genetic crosses with the male gamete-defective lines did not restore the KO phenotype. **(C)** Quantification of oocysts on day 15. Sixty-two midguts from each group were dissected and observed. Significant differences were observed in WT vs WT-mCherry×Alp5b KO (**P=0.0038), WT vs WT-GFP×Alp5a KO (**P=0.0019), WT vs Pb47 KO (****P<0.0001), WT vs Pb45/48 KO (****P<0.0001), WT vs Pb47 KO×Alp5b KO (**P=0.0071), WT vs Pb48/45 KO×Alp5b KO (****P<0.0001), WT vs Pb47 KO×Alp5a KO (**P=0.0050), and WT vs Pb48/45 KO×Alp5a KO (****P<0.0001). **(D)** Oocyst area on day 15 post-infection. A total of 210 oocyst areas were quantified in each group. Significant differences were observed in WT vs WT-mCherry×Alp5b KO (*P=0.0377), WT vs WT-GFP×Alp5a KO (*P=0.0203), WT vs Pb47 KO×Alp5b KO (*P=0.0334), WT vs Pb48/45 KO×Alp5b KO (****P<0.0001), WT vs Pb47 KO× Alp5a KO (*P=0.0435), and WT vs Pb48/45 KO× Alp5a KO (****P<0.0001). Unpaired Student’s t test was used to determine the statistical significance. The data were pooled from three independent experiments and are presented as the mean ± SEM.

### Alp5s regulate DNA segregation during male gametogenesis

The male gametocyte undergoes three rapid rounds of DNA division during male gametogony, reaching an 8 N state. After division, this DNA distributed to the eight flagellar microgametes coincided with axoneme formation. To determine the defect in male gametogony, activated gametocytes were immunostained with an anti-tubulin antibody to identify activated male gametes and free microgamete. The diffuse signal of a single nucleated cell was considered a female gametocyte/macrogamete on the same slide. We found no differences in the number of activated male or female gametes, but there was a difference in the pattern of DNA allocation in activated gametocytes. The DNA distribution was homogenous, flagellar and condensed in the WT parasites, with enlarged activated gametocyte nuclei. However, the DNA was residual and irregularly distributed in the Alp5 KO parasites (Figure 5A). The percentage of irregularly distributed Alp5a KO and Alp5b KO cells was greater than that of WT cells (Figure 5B). We found no difference in the DNA content of activated male gametocytes (enlarged nuclei coinciding with axonemes) or female gametocytes/gametes (single nuclei with diffuse tubulin signals). There was a significant reduction in the DNA content in the male gamete (Figure 5C and D). The DNA was quantified via the CTCF method via ImageJ software. The reduced DNA content in the male gametes of Alp5 KO parasites did not affect the morphology of zygotes or ookinetes. Next, we checked the DNA content in zygotes and ookinetes, which was significantly reduced in Alp5 KO parasites (Figure 5 E, F and G). The reduced DNA content in the ookinete did not affect its gliding ability (Figure S9C). Next, we analyzed the DNA in the oocysts on different days via Hoechst staining. These oocysts were imaged, and the DNA content was quantified via Fijji software. The DNA content in the KO oocysts was negligible because of impaired development. However, the DNA contents in the WT and complemented lines were similar (Figure 5H and I). The DNA content in the oocysts of the KO and female gamete-defective crossed lines was comparable to that of the WT, whereas there was no change in the DNA content in the KO lines crossed with the male gamete-defective line (Figure 5J and K). These results suggest that Alp5s are essential for DNA segregation during male gametogenesis. Alp5 KO parasites undergo meiosis and ookinete formation despite lower DNA content but are arrested at the oocyst stage.

**Figure 5.**
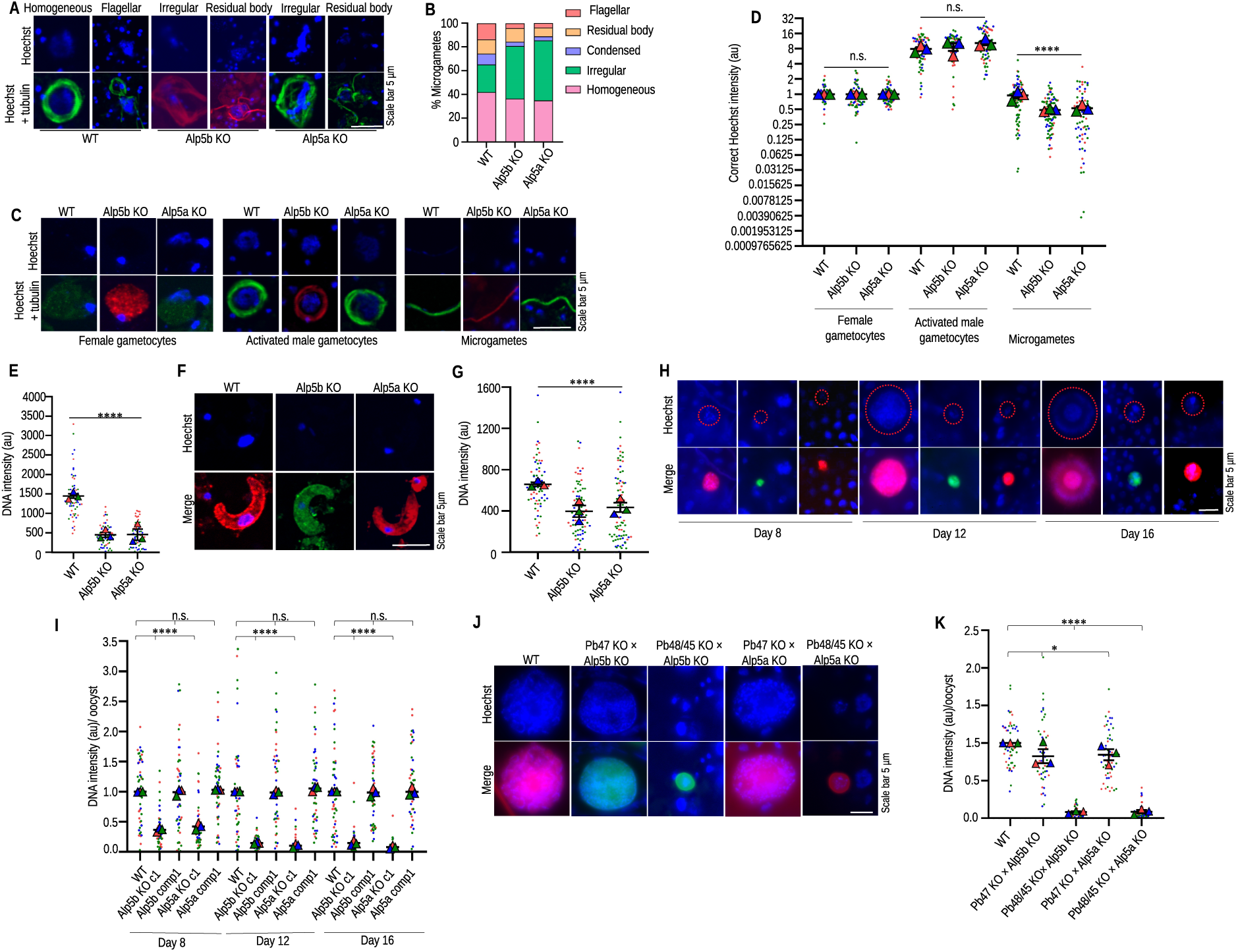
Impaired DNA segregation in Alp5 KO parasites. **(A)** Gametocytes were activated and fixed at 15 mpa. DNA was stained with Hoechst, and an anti-tubulin antibody was used to identify activated male gametocytes. **(B)** Different forms of microgametes identified on the basis of quantification of the DNA localization pattern in activated male gametocytes. **(C)** IFA images of female gametocytes, activated male gametocytes and microgametes. Activated male gametocytes and microgametes were identified by anti-tubulin antibody staining. The female gametocytes were identified via diffuse tubulin staining. DNA was stained with Hoechst. **(D)** DNA content of activated male and female gametocytes and microgametes. The DNA content was normalized to the mean DNA content of female gametocytes from the same slide. The data were comparable between WT and Alp5 KO strains in terms of the number of female gametocytes (P=0.3126) and activated male gametocytes (P=0.0842). The DNA content significantly differed between WT and Alp5 KO microgametes (****P<0.0001). The Kruskal‒Wallis test was used to determine statistical significance. **(E)** Quantification of the zygote DNA content (a.u., arbitrary unit). We detected significant differences among the WT, Alp5a KO and Alp5b KO parasites (****P<0.0001). The DNA content of 60 zygotes per group was analyzed. **(F)** Compared with WT ookinetes, Alp5 KO ookinetes presented a lower DNA content. **(G)** Quantification of ookinete DNA content (a.u., arbitrary unit). We detected significant differences between WT and Alp5 KO parasites (****P<0.0001). The DNA contents of 70 (WT), 83 (Alp5b), and 83 (Alp5a) ookinetes were quantified. **(H)** Hoechst-stained oocysts showing the DNA content on different days. **(I)** The average DNA intensity of 50 oocysts was significantly different between WT and Alp5 KO parasites on days 8 (****P<0.0001), 12 (****P<0.0001) and 16 (****P<0.0001). The DNA content was comparable between WT, ALP5b comp and ALP5a comp on day 8 (P=0.7954), day 12 (P=0.6828), and day 16 (P=0.9362). The statistical significance was analyzed via the Kruskal‒Wallis test**. (J)** Hoechst-stained oocysts showing the DNA content after the genetic cross. The DNA content was restored in Alp5 KO parasites crossed with female gamete-defective lines. Genetic crosses with male gamete-defective lines did not affect the oocyst DNA content. **(K)** The DNA content of 50 oocysts in Alp5 KO parasites was quantified and normalized to the mean DNA intensity of WT oocysts. A significant difference was observed in WT vs Pb47 KO×Alp5b KO (*P=0.0330), WT vs Pb48/45 KO×Alp5b KO (****P<0.0001), WT vs Pb47 KO×Alp5a KO (*P=0.0172), and WT vs Pb48/45 KO×Alp5a KO (****P<0.0001). Statistical analysis was performed via an unpaired Student’s t test.

## Discussion

The actin cytoskeleton is involved in a plethora of cellular functions (46). The rate-limiting step in actin filament assembly is the nucleation step. One of the major actin-nucleating factors in cells is the seven subunit Arp2/3 complex. The Arp2/3 complex is highly conserved in all eukaryotes (32, 33, 36). In this study, we investigated the functions of the human Arp2- and Arp3-related *Plasmodium* proteins Alp5a and Alp5b, which revealed that these proteins have distinct cellular functions in DNA segregation during male gametogenesis. The STRING database indicated the presence of five subunits of Arp2/3 complex proteins in *Plasmodium*. A study recently identified a non-canonical Arp2/3 complex in *Plasmodium* that consists of only five subunits (47). We found that the Arp2/3 complex subunits, Alp5a and Alp5b, localize from the nucleus to the axoneme and remain in the residual body after exflagellation. Parasites lacking Alp5, form subhaploid male gametes that produce functional zygotes and motile ookinetes. However, Alp5 KO parasites are arrested in a delayed-death-like manner at the oocyst stage, leading to complete transmission block within mosquitoes (Figure 6).

**Figure 6.**
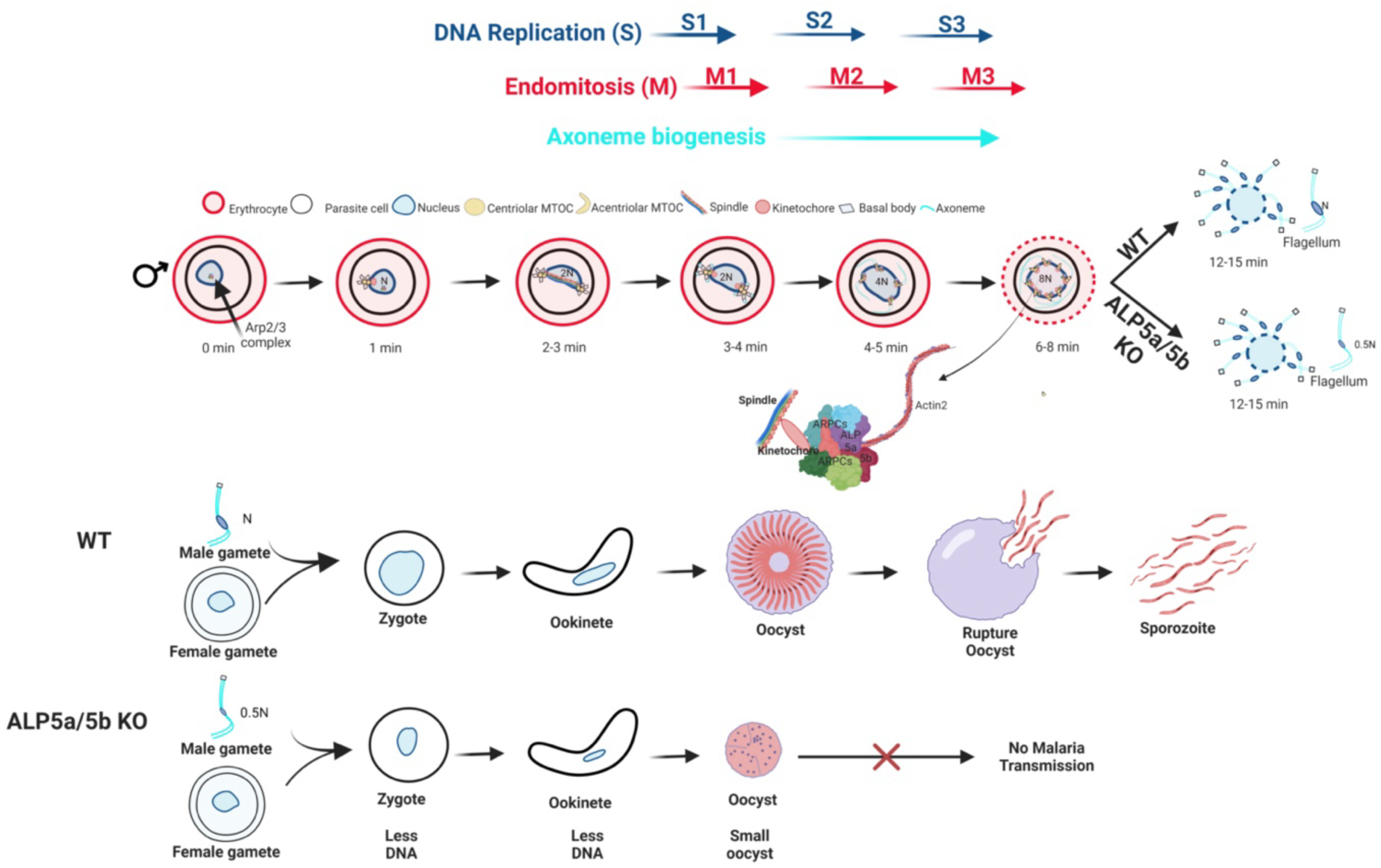
Model of non-canonical Arp2/3 complex-mediated nucleation of actin filaments. During male gametogony, actin stabilizes the endomitotic spindle assembly. The parasites lacking Alp5 show reduced DNA content from the male gamete to the oocyst. The reduced DNA content in the male genome could cause a defect in DNA replication that affects oocyst development.

Alp5s are predominantly expressed in the nucleus and are localized from the nucleus to the axoneme. We found that Alp5s colocalize with F-actin and that Alp5 KO parasites exhibit defects in DNA segregation and male gamete fertility. Like humans, Arp3, Arp2, Alp5a and Alp5b interact with each other. Our results indicate that, like canonical Arp2/3 complexes, *Plasmodium* Arp2/3 preferentially nucleates actin filaments. It has been reported that actin 2 is present in male gametocytes, female gametes, and zygotes (25, 48) and that it is associated with the nucleus in the male gametocyte and in the zygote. Although actin 2 is predominantly found in the nucleus, its signals are also detected in the cytoplasm (23). Similarly, we detected the Alp5s signal in the cytoplasm, indicating its additional role beyond nuclear function. Surprisingly, the indispensability of Alp5a and Alp5b simultaneously and the inhibition of *P. berghei* and *P. falciparum* blood stage development with an Arp2/3 complex inhibitor suggest that the complex does play a role during blood stage development. The similar F-actin/G-actin ratios in WT and Alp5 KO parasites support the dispensability of individual Alp5 subunits. We speculate that parasites lacking one Alp5 subunit can nucleate actin. However, parasites that lack both Alp5 subunits or are inhibited by Arp2/3 complex inhibitors fail to nucleate actin. The STRING database revealed the interaction of *P. berghei* Alp5 with Formin 2. We hypothesize that the *Plasmodium* Arp2/3 complex lacking one subunit is possibly stabilized by Formin 2, or that they nucleate actin together. The malaria parasite Formin regulates actin polymerization, and the parasite uses polymerized actin throughout its lifecycle. Formin 1 is essential for invading merozoites, and Formin 2 is critical for efficient cell division in *P. falciparum* (29). Formin inhibitors block multiple asexual and sexual parasite stages of development (29). On the basis of the cytosolic localization of Formin 2 (29), it is tempting to speculate that Formin 2, together with the Arp2/3 complex, is involved in the nucleation of actin 1. However, together with previous reports (49), our results indicate that Formin 2 is the primary cytosolic F-actin nucleator and that the Arp2/3 complex plays a supporting role during *Plasmodium* asexual blood stage development. Malaria parasites encode two actin isoforms, actin 1 and actin 2. Actin 1 is indispensable during blood stage development (22, 50, 51). Actin 2 is dispensable in blood stage but has essential functions in male gametogenesis and oocyst development (25, 26). Actin 2 mainly localizes to the nuclear spindle and, to some extent, to the cytoplasm and actin 2 mutant male gametocytes fail to exflagellate (23, 26). Attempts to complement actin 2 function with actin 1 partially restored male gametogenesis, resulting in the formation of oocysts, but these oocysts failed to form sporozoites (25). The inability of actin 1 to complement the function of actin 2 suggests that the Arp2/3 complex nucleates actin 2 without associating with Formin 2.

We found defects in male gamete integrity in Alp5 KO parasites via genetic cross-experiments. The sporogony of the restored oocysts was normal, and further sporozoites infect the salivary glands and mammalian hosts normally. The lack of Alp5 expression in the sporozoites and liver stage indicates that the Arp2/3 complex plays no role during these stages. Therefore, the primary function of the Arp2/3 complex is likely the nucleation of actin 2. We found that Alp5 KO male gametes contained only half of the genome. The localization of Alp5s to the axoneme suggests that Arp2/3 act at the level of DNA segregation. A recent report revealed that the Arp2/3 subunit ARPC1 localizes to the mitotic spindle, interacts with the kinetochore protein AKiT7, and acts at the kinetochore-spindle interphase (47). ARPC1 KO parasites fail to form and maintain the eight kinetochore foci during DNA condensation and subsequent exflagellation (47). Together with these results, we speculate that actin polymerization is required to stabilize the attachment of kinetochores to the spindle during axonemal movement. In Alp5 KO parasites, a greater proportion of irregular microgametes indicates a defect in DNA segregation.

The Alp5 KO parasites formed oocysts but did not undergo sporogony and presented a delayed death phenotype. *P. berghei* lacking end-binding protein 1 (EB1) (52–54) resembles the phenotype of Alp5 KO parasites (52). EB1 connects the kinetochores to the spindle during male gametogenesis, and EB1 KO parasites present a similar delayed death phenotype (52). Early oocyst phenotypes have also been reported for mutant parasites lacking the formin-like protein MISFIT (55), aurora-related kinase-2 (Ark2) (52), and the microtubule motor kinesin-8X (56). The Alp5 KO parasites with a defect in the male gamete with dying oocysts contain a complete genome provided by the female gamete. The reduced DNA content in the male genome could cause a defect in DNA replication that affects oocyst development. Whether this phenotype is due to male gamete defects or whether the Arp2/3 complex has an independent role during oocyst development needs further investigation.

In conclusion, we show here that *Plasmodium* Alp5s are essential for malaria parasite transmission to mosquito. Arp2/3 complex plays a supporting role during blood stage development for actin 1 polymerization, whereas its role in actin 2 nucleation is indispensable during male gametogenesis. Further work is needed to fully understand the role of the Arp2/3 complex and the mechanisms of cell division in the male gamete. Taken together, our results show that Alp5 plays an essential role in male gametocyte development and interfering with its function may disrupt the cycle of malaria transmission.

## Materials and methods

### Bioinformatics analysis

The protein sequences of *Plasmodium berghei* Alp5a and Alp5b were retrieved from PlasmoDB (https://plasmodb.org). The amino acid sequences were queried via NCBI protein BLAST (P BLAST) against model organisms. Owing to the unavailability of experimentally determined structures, the protein structures of PbAlp5a and PbAlp5b were predicted via AlphaFold 2 (https://colab.research.google.com/github/sokrypton/ColabFold/blob/main/AlphaFold 2.ipyn). The quality of the predicted structure was assessed via SAVES v6.0 (https://saves.mbi.ucla.edu/). To assess structural similarity and protein folding, the structure of Alp5a was superimposed onto the known human Arp3 protein structure, and Alp5b was superimposed onto the human Arp2 structure (human Arp2:6YW6 and Arp3:6UHC proteins from the PDB database https://www.rcsb.org/). The amino acid sequences of Alp5a and Alp5b were submitted to the STRING database to investigate potential interactions between these proteins. Further protein-protein interactions were predicted via the ClusPro protein-protein interaction server.

### Parasites, mosquitoes and mice

*Plasmodium berghei* ANKA (MRA 311) and *P. berghei* ANKA GFP (MRA 867 507 m6cl1) were obtained from BEI Resources, USA. *Anopheles stephensi* mosquitoes were reared in a walk-in environmental chamber maintained at 28°C with 80% relative humidity and 12 h light and dark cycles. Mosquitoes were fed with 10% sugar syrup soaked in cotton pads. Female *Anopheles* mosquitos were allowed to probe for a blood meal on Swiss mice for egg laying. The parasites were transmitted to mosquitoes as described previously (57). The walk-in environmental chamber was maintained at 19°C with 80% relative humidity and 12 h light and dark cycles to keep the infected mosquitoes. Swiss albino and C57BL/6 mice (6-8 weeks old) were used in this study. All animal experiments were performed according to the Institutional Animal Ethics Committee guidelines at CSIR-Central Drug Research Institute, India (IAEC reference no: IAEC/2018/3 to IAEC/2023/15). Chloroquine-sensitive *P. falciparum,* strain 3D7 (MRA-151, BEI Resources, USA) was used to assess the effects of the inhibitors. Parasites were cultured in human RBCs (approval #CDRI / IEC / 2017 / A4 by the Institutional Ethics Committee-Human Research) at 2% haematocrit in HEPES-modified RPMI-1640 medium (Sigma Aldrich, USA) supplemented with 1% glucose, 0.2% sodium bicarbonate, 100 µM hypoxanthine (Sigma-Aldrich #Cat. H9636), gentamycin (Gibco #Cat. 15750-060), (25 μg/ml) and 0.5% Albumax II (Gibco #Cat. 11021-037). The parasites were regularly synchronized with 5% D-sorbitol (58). Human liver hepatocellular carcinoma (HepG2) cells (ATCC) were cultured in DMEM (Sigma-Aldrich #Cat. D5648) enriched with 10% FBS (Biological Industries #Cat. 04-121-1A), 0.2% NaHCO_3_ (Sigma-Aldrich #Cat. S5761), 1% sodium pyruvate (Genetix #Cat. CC4016), and 1% penicillin‒streptomycin (Gibco #Cat. 15140-122) maintained at 37°C with 5% CO2.

### Generation of Alp5b knockout and complemented parasite lines

To delete the Alp5b gene, the wild-type locus was replaced with a targeting cassette via double crossover homologous recombination. Two fragments, F1 (0.545 kb) and F2 (0.6 kb), were amplified from the 5’ and 3’ UTR regions via the primer sets 1611/1612 and 1613/1614, respectively. Fragments F1 and F2 were subsequently cloned into the pBC-GFP-hDHFR-yFCU vector at *Sal*I and *Not*I/*Asc*I, respectively. The final targeting cassette was separated from the vector backbone via digestion with *Xho*I/*Asc*I and transfected into *P. berghei* ANKA schizonts as described previously (59). Drug-resistant parasites that appeared after pyrimethamine selection were observed for GFP fluorescence. The genomic DNA was isolated from the GFP expressing drug-resistant parasites, and correct 5’ and 3’ site-specific integration was confirmed by PCR via the primer sets 1665/1225 and 1666/1215, respectively. The clonal lines were obtained by limiting dilution of the parasites and injecting them into Swiss mice. The clonal KO lines were confirmed via the primers 1671/1672. The Alp5b complemented line was generated by reintroducing the ORF via double crossover homologous recombination. A fragment encompassing the 5’UTR, ORF and 3’UTR was amplified via the primer set 1611/1614 and transfected into Alp5b KO schizonts. The transfected parasites were negatively selected with 5-fluorocytosine (Sigma-Aldrich #Cat. F7129) as previously described (60). Gene restoration was confirmed by PCR via the primers 1671/1672.

### Generation of Alp5a knockout and complemented parasite lines

To disrupt the Alp5a locus, two fragments, F3 (0.59 kb) and F4 (0.52 kb), were amplified via the primer sets 1895/1896 and 1897/1898, respectively. Fragments F3 and F4 were sequentially cloned into a pBC-mCherry-tgDHFR vector at *Kpn*I/*Cla*I and *Not*I/*Asc*I, respectively. The plasmid was linearized via *Kpn*I/*Asc*I and transfected into *P. berghei* ANKA schizonts. Genomic DNA was isolated from mCherry expressing pyrimethamine-resistant parasites, and 5’ and 3’ site-specific integration was confirmed by diagnostic PCR via primers 2089/1225 and 1913/2090, respectively. Clonal lines were obtained as described above and confirmed via the primers 2091/2092. To restore gene function, fragment F5 encompassing the Alp5a-expressing cassette (5’UTR+ORF+3’UTR) was amplified via primers 2268/2269 and subsequently cloned into pBC-P230-mCherry-hDHFR at *Cla*I and *EcoR*I. The linearized cassette was transfected into Alp5a KO schizonts, which were selected via the WR99210 drug (Sigma-Aldrich #Cat. SML2976) as previously described (61). Correct 5’ and 3’ site-specific integration was confirmed by diagnostic PCR via primer sets 2270/2271 and 1215/2272, respectively. The restoration of the ORF was confirmed via the primer set 2091/2092. The primer sequences are given in Table S1.

### Attempts to generate Alp5a/Alp5b double knockout parasites

Two strategies were employed to generate Alp5a/Alp5b double KO parasites. In the first strategy, the linearized cassette of Alp5b was transfected into Alp5a KO schizonts, which were selected with the WR99210 drug. In the second strategy, Alp5b and ALP5a targeting cassettes were cotransfected into *P. berghei* ANKA schizonts, which were selected with pyrimethamine (Sigma-Aldrich #Cat. 46706).

### Generation of Alp5a and Alp5b transgenic parasites

For Alp5b tagging with 3XHA-mCherry, two fragments, F6 (0.54 kb) and F7 (0.60 kb), were amplified via the primer sets 1675/1676 and 1677/1614 and cloned into the pBC-3XHA-mCherry-hDHFR vector at *Xho*I/*Bgl*II and *Not*I/*Asc*I, respectively. For Alp5a tagging with 3XHA, two fragments, F8 (0.61 kb) and F4 (0.52 kb), were amplified via the primers 2128/2129 and 1897/1898 and cloned into the pBC-3XHA-hDHFR vector at *Xho*I/*Bgl*II and *Not*I/*Asc*I, respectively. The vector was linearized via *Xho*I/*Asc*I and transfected into *P. berghei* ANKA schizonts as described above. Correct 5’ and 3’ site-specific integration was confirmed by diagnostic PCR via the primers 1724/1392 and 1215/1666 for Alp5b and 2130/1218 and 1215/2090 for Alp5a. The primer sequences are given in Table S1.

### Generation of gamete-defective lines

The 6-cysteine proteins, P47 (PBANKA_1359700) and P48/45 (PBANKA_1359600) targeting constructs for generating female and male gamete-defective lines, respectively, were obtained from the PlasmoGEM resource (https://plasmogem.umu.se/pbgem/) (62). The plasmids were linearized via *Not*I, transfected into *P. berghei* ANKA schizonts and selected with pyrimethamine. Genomic DNA was isolated from drug-resistant parasites, and site-specific integration of the P47 and P48/45 targeting cassettes was confirmed by diagnostic PCR viaprimers 2232/2035 and 2230/2231, respectively. Clonal lines were obtained by limiting dilution of the parasites, and the absence of P47 and P48/45 ORFs was confirmed via primers 2233/2234 and 2230/2231, respectively. The primer sequences are given in Table S1.

### Asexual blood-stage propagation and gametocyte development

Two groups of Swiss mice (5 mice/group) were injected intravenously with 200 µl of blood containing 0.5% parasitemia. Blood growth and gametocyte development were monitored via Giemsa-stained blood smears.

### Analysis of parasite development in mosquitoes

Swiss mice were injected with WT or KO parasites as described above. The mice positive for gametocytes were anesthetized and placed in starved mosquito cages for a blood meal. To analyze the different developmental stages of parasites, mosquitoes were dissected under a stereo-zoom microscope. Blood boluses were collected 10– 20 min or 18–22 h postfeeding to analyze gametes and ookinetes, respectively, and slides were prepared for live imaging as previously described (60). The blood boluses was crushed and diluted, and the ookinetes were counted via a hemocytometer. Mosquito midguts were collected on days 8-18 to determine the oocyst numbers and areas. The oocyst area was determined via Fiji software. Midgut and salivary gland sporozoites were enumerated on days 14 and 19-22 post-blood meal, respectively, as previously described (61).

### Genetic cross

A genetic cross between the mutant and WT lines was performed. For this purpose, blood was collected from the mice infected with WT or KO parasites and mixed at a ratio of 1:1. Two hundred microliters of blood with 0.5% parasitemia was injected into a group of Swiss mice. Mosquitoes were allowed to probe for a blood meal in infected mice. Mosquitoes were dissected on days 15 and 19-22 after a blood meal to observe oocysts and salivary gland sporozoites, respectively.

### Gametocyte purification and activation

Gametocytes were purified via a nycodenz gradient. Swiss mice were treated with phenylhydrazine (6 mg/ml) (Sigma-Aldrich #Cat. 114715) for 3 days and then intraperitoneally injected with 200 µl of blood containing 1 % parasitemia. After 3–4 days, the presence of gametocytes was confirmed by Giemsa-stained blood smears. Blood was collected in RPMI, loaded on a 50% nycodenz gradient, and centrifuged for 20 minutes at 200 × g. The gametocyte ring that appeared on the interphase was collected and washed 2–3 times with PBS. The purified gametocytes were activated using an exflaggelation medium containing 25 mM HEPES (Sigma-Aldrich #Cat. 83264) and 100 µM xanthurenic acid (Sigma-Aldrich #Cat. D120804) supplemented with RPMI medium (Sigma-Aldrich #Cat. 23400-013) at pH 8.0. Different stages were observed, either live or after the spot was fixed with 4% PFA (Sigma-Aldrich #Cat. HT5012).

### Exflagellation centres

An exflagellation assay was performed by incubating 10 µl of purified gametocytes in 40 µl of exflagellation medium for 5 –10 minutes at 20°C. The exflagellation centres were observed under a phase contrast microscope at 100x magnification and counted per field.

### Ookinete motility assay

Blood boluses were isolated, diluted, and mixed with an equal volume of Matrigel (BD Biosciences #Cat.356231). A 5 µl drop was placed on a slide, gently covered with a coverslip, and then incubated for 20 minutes at RT. After incubation, the slides were imaged under a Nikon Eclipse 80i microscope at 100x magnification, and a time-lapse video of 61 loops for 5 minutes was recorded.

### Generation of the antibodies

Antibodies against Alp5b and P25 were generated in rats. KLH-conjugated peptides of Alp5b (Cys-SNKKKWITKNEYSANP) (GLS #Cat. 998808). and P25 (SPNTQCKNGFLAQMSN-Cys) (GLS #Cat.997976) were synthesized by GL Biochem. The peptides were immunized once with complete Freund’s adjuvant and twice with incomplete Freund’s adjuvant. The rats were bled, and the serum was collected for further analysis.

### In vivo infectivity of sporozoites

Infected mosquitoes were used for sporozoite transmission 22 days post-blood meal, as previously described (63). Briefly, C57BL/6 mice were anesthetized and kept on the mosquito cages. Fifty mosquitoes per C57BL/6 mouse were used. Infection was observed via Giemsa-stained blood smears.

### In vitro development of exoerythrocytic stages

To observe the development of the parasite in hepatocytes, salivary gland sporozoites were added to HepG2 cells as previously described (64). Briefly, HepG2 cells were seeded in a 48-well plate (55,000 cells/well) containing collagen-coated 9 mm glass coverslips. The next day, salivary gland sporozoites (5,000/well) were added to the wells, and the plate was spun at 310 × g for 4 minutes. The infected culture was kept in a CO_2_ incubator set at 37°C with 5% CO_2_. The medium was changed every 12 h, and the culture was fixed with 4% paraformaldehyde for 20 minutes at RT.

### Immunofluorescence assay (IFA)

Parasites at different stages were harvested, and IFA was performed as previously described (65). Briefly, parasites were fixed for 20 minutes at RT with 4% paraformaldehyde. Blood smears and sporozoite slides were permeabilized with 0.1% Triton X-100 (Sigma‒Aldrich #Cat. T8787) for 10 minutes at RT, and liver stage samples were permeabilized with chilled methanol. The samples were blocked with 1% BSA for 1 h at RT and incubated with primary antibodies for 1 h at RT or overnight at 4°C. The primary antibodies used were anti-CSP (diluted 1:1000), anti-HA (diluted 1:1000), anti-actin (diluted 1:1000), anti-g377 (diluted 1:100) (60), anti-tubulin (diluted 1:100), anti-P25 (diluted 1:100), and anti-MSP1 (diluted 1:5,000). The following secondary antibodies were used: Alexa Fluor 488-conjugated anti-rabbit IgG (diluted 1:1,000), (Invitrogen #Cat. A11008), Alexa Fluor 594-conjugated anti-rabbit IgG (diluted 1:1,000), (Invitrogen #Cat. A21442), Alexa Fluor 594-conjugated anti-rat IgG (diluted 1:1,000), (Invitrogen #Cat. A11007), Alexa Fluor 488-conjugated anti-rat IgG (diluted 1:1,000), (Invitrogen #Cat. A11006), Alexa Fluor 488-conjugated anti-mouse IgG (diluted 1:1,000), (Invitrogen #Cat. A11001) and Alexa Fluor 594-conjugated anti-mouse IgG (diluted 1:1,000), (Invitrogen #Cat. A11032). Nuclei were stained with Hoechst 33342 (Invitrogen #Cat. 62249), and samples were mounted with Prolong Diamond antifade reagent (Invitrogen #Cat. P36970). Images were captured via a 100x oil immersion lens under a Nikon Eclipse 80i fluorescence microscope or Leica DM 3000 LED microscope.

### Analysis of DNA content

The developing male gametocytes were identified by immunostaining with an anti-tubulin antibody, and diffuse signals in female gametocytes were used to differentiate them. The ookinetes were immunostained with an anti-P25 antibody. For DNA content analysis of the oocysts, the midgut was isolated at 8, 12, 15, and 16 days post-feeding and fixed with 4% PFA for 30 minutes at RT. The fixed midguts were washed twice with cold PBS for 5 minutes, permeabilized and immunostained with an anti-CSP antibody. Nuclei were stained with Hoechst 33342 and mounted with ProLong Diamond antifade reagent. Hoechst-stained DNA was quantified via Fiji software. The total fluorescence intensity was measured after subtracting the background signal. The intensity was normalized to that of female gametocytes or single-nucleated parasites. The software (https://theolb.readthedocs.io/en/latest/imaging/measuring-cell-fluorescence-using-imagej.html) was used to measure the fluorescence intensity.

### Determination of the G-actin/F-actin ratio

The G-actin/F-actin ratio was determined via a G-actin/F-actin in vivo assay kit (Cytoskeleton, Inc. #Cat. BK037) according to the manufacturer’s instructions. The blood was lysed with saponin, and the parasite pellet was washed four times with 1X PBS, resuspended in LAS2 buffer, and incubated at 37°C for 10 minutes. The samples were centrifuged at 20,913 × g for 5 min and then transferred to a fresh tube for a second spin at 100,000 × g for 1 h in an ultracentrifugation. The supernatant and pellet were separated and lysed in Laemmli buffer. The samples were resolved via 10% SDS‒PAGE and analyzed via western blotting.

### Effects of inhibitors on erythrocytic schizogony

Synchronized *P. falciparum* parasites were incubated with different concentrations of cytochalasin-D (Sigma-Aldrich #Cat. C8273), jasplankinolide (Sigma-Aldrich #Cat. 420127), and the Arp2/3 complex inhibitor, CK-666 (Sigma-Aldrich #Cat. 182515) throughout the culture. A schizont culture was set up as described above to examine the effects of the inhibitors on the asexual blood stages of *P. berghei*. The inhibitors were added to the culture and incubated at 37°C for 22–24 h. The growth and morphology of the parasites were monitored via Giemsa-stained smears.

### Immunoprecipitation

Purified gametocytes of the Alp5a-3XHA and WT strains were washed four times with DPBS and crosslinked with 5 mM DSP (dithiobis succinimidyl propionate; Thermo Scientific #Cat. 22586). The DSP-crosslinked samples were incubated for 30 minutes in the dark and centrifuged at 20,913 × g for 5 minutes. The pellet was lysed on ice in 500 µl of RIPA buffer for 30 minutes. The mixture was centrifuged at 20,913 × g for 15 minutes, and the supernatant was transferred to a fresh microcentrifuge tube. A total of 25 µl of anti-HA magnetic beads (Thermo Scientific #Cat. 88838X) was washed with 0.05% TBST four times in a magnetic stand (Stem Cell Technologies #Cat. 18000). The beads were incubated with parasite lysate for 30 min at RT on a tube rotator. After incubation, the beads were washed with TBST followed by ultrapure water, eluted in 2x Laemmli buffer (Bio-Rad #Cat. 1610737), and boiled for 5 minutes. The samples were resolved via 10% SDS‒PAGE, transferred to nitrocellulose membranes, and probed with anti-Alp5b and anti-HA antibodies.

### Western blotting

Western blotting was performed as previously described (65). Briefly, samples were resolved via 10% SDS‒PAGE and transferred onto a nitrocellulose membrane (Bio-Rad #Cat. 1620112). The membrane was blocked with 1% BSA/PBS and incubated with anti-HA (diluted 1:1,000), (CST #Cat. C29F4) or anti-Hsp70 (diluted 1:1,000) antibodies. The blot was washed 3–4 times with 1X PBST and then incubated with HRP-conjugated anti-rabbit (diluted 1:5,000) (Amersham Biosciences #Cat. NA934V) or anti-mouse IgG (diluted 1:5,000) (Amersham Biosciences #Cat. NA931V). The signals were detected via an enhanced chemiluminescence (ECL) substrate (Bio-Rad, #Cat. 170--5060) and imaged via a ChemiDoc XRS+ system (Bio-Rad, USA).

### Software and statistics

All the statistical analyses were performed via GraphPad Prism 9 software. More than two groups were analyzed via one-way ANOVA, and two groups were compared via Student’s t test. *P<0.05, **P<0.01, ***P<0.001, ****P<0.0001, n.s.; not significant.

## Data and Materials Availability

All the data are available within this manuscript and the raw data are available from the corresponding author upon reasonable request. Materials generated in this study are available from the corresponding author upon request.

## Acknowledgments

We thank BEI Resources, USA, for the parasite strains and plasmids and Dr. Robert Menard and Dr. Kota Arun Kumar for the KO and tagging vectors. We thank Dr. Anthony A. Holder for MSP1 antibodies. We thank Dr. Pratik Narain Srivastava for helping us analyze the in silico data and Dr. Shabeer Ali for the *P. falciparum* drug assay. We acknowledge the CSIR-CDRI Intravital microscopy facility. The Council of Scientific and Industrial Research, Department of Biotechnology, and University Grants Commission Government of India research fellowships supported AV, EP and N. SM acknowledges grants from the Indian Council of Medical Research (6/9-7(306)/2022/ECD-II) and the Science and Engineering Research Board (CRG/2022/003848).

## Notes

### Competing Interest Statement

The authors have declared no competing interest.

