## Supplementary material for "*Plasmodium* actin-like proteins are essential for DNA segregation during male gametogenesis and malaria transmission"

Satish Mishra

**This PDF file includes:**

Figures S1 to S13

Tables S1

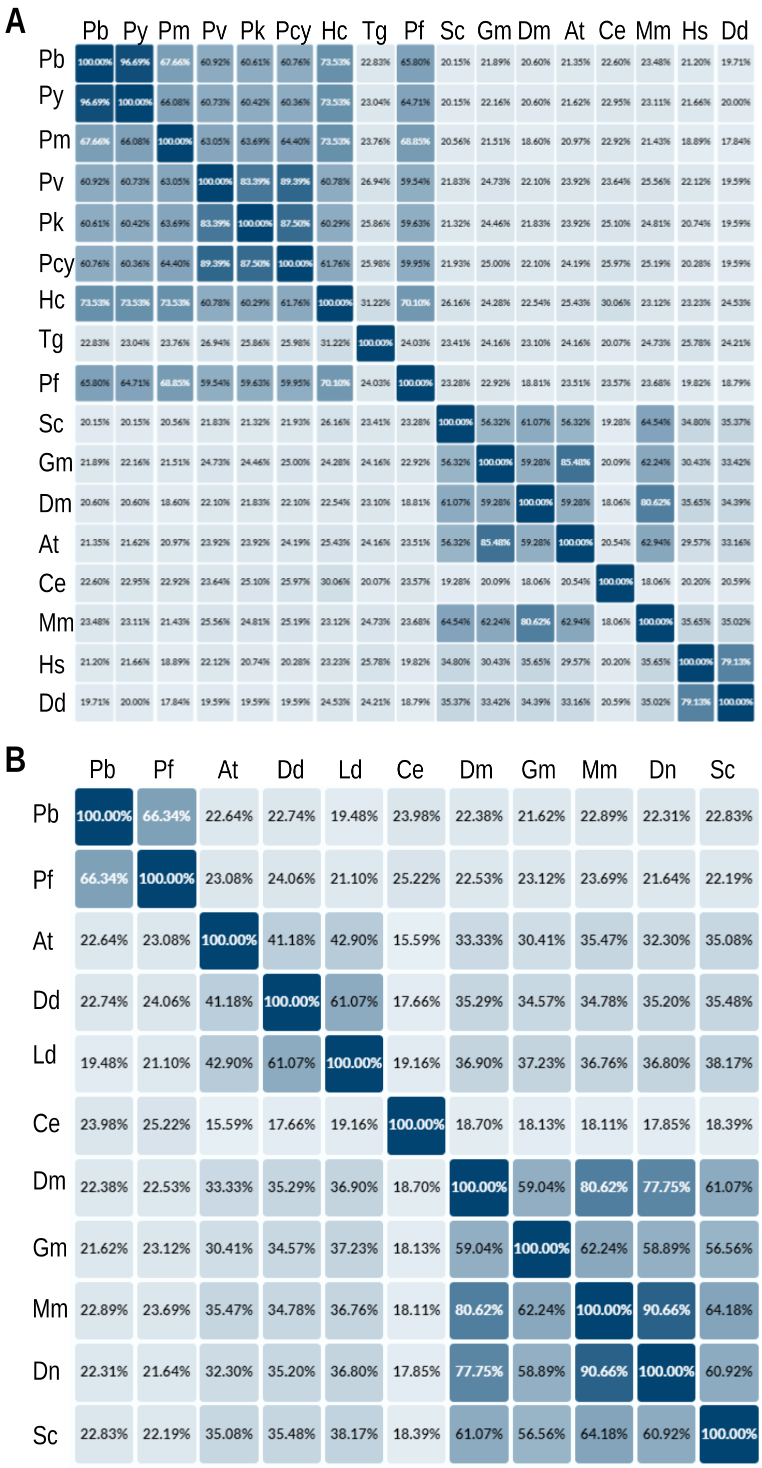


**Figure S1. In silico amino acid analysis of ALP5a and ALP5b. (A)** The % sequence similarity matrix of Alp5a with *Plasmodium* and other model organisms. **(B)** The % sequence similarity matrix of Alp5b. The full initials are as follows: *Pb- Plasmodium berghei, Py- Plasmodium yoelii, Pm- Plasmodium malariae, Pv- Plasmodium vivax, Pk- Plasmodium knowlesi, Pcy- Plasmodium cynomolgi, Hc- Hepatocystis, Tg- Toxoplasma gondii, Pf- Plasmodium falciparum, Sc-Saccharomyces cerevisiae, Gm- Glycine max, Dm- Drosophila melanogaster, At- Arabidopsis thaliana, Ce- Caenorhabditis elegans, Mm- Mus musculus, Hs- Homo sapiens* and *Dd- Dictyostelium discoideum, Ld- Leishmania donovani* and *Dn- Danio.*

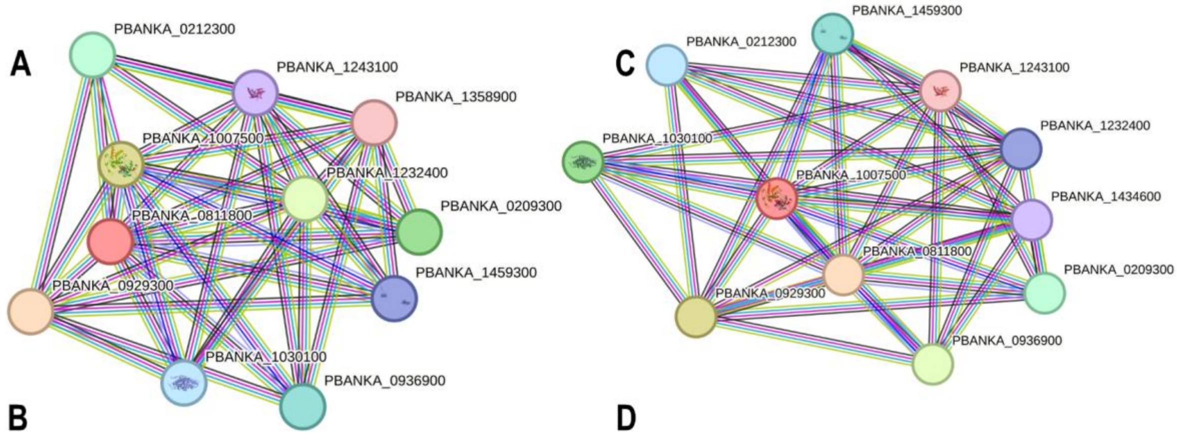


**B**

| Gene name | Predicted partners of Alp5a (PBANKA_0811800) |
|---|---|
| PBANKA_0212300 | actin-like protein, putative |
| PBANKA_1243100 | F-actin-capping protein subunit Alpha |
| PBANKA_1358900 | dynactin subunit 2, putative |
| PBANKA_1007500 | actin-like protein, putative Alp5b |
| PBANKA_1232400 | F-actin-capping protein subunit beta |
| PBANKA_0209300 | actin-related protein ARP1 |
| PBANKA_1459300 | actin I |
| PBANKA_0929300 | actin-related protein 2/3 complex subunit 1, putative |
| PBANKA_1030100 | actin II |
| PBANKA_0936900 | actin-like protein, putative |
| PBANKA_1434600 | formin 2, putative |

**D**

| Gene name | Predicted partners of Alp5b (**PBANKA_1007500)** |
|---|---|
| PBANKA_0212300 | actin-like protein, putative |
| PBANKA_1243100 | F-actin-capping protein subunit alpha |
| PBANKA_1358900 | dynactin subunit 2, putative |
| PBANKA_0811800 | actin-like protein, putative Alp5a |
| PBANKA_1232400 | F-actin-capping protein subunit beta |
| PBANKA_0209300 | actin-related protein ARP1 |
| PBANKA_1459300 | actin I |
| PBANKA_0929300 | actin-related protein 2/3 complex subunit 1, putative |
| PBANKA_1030100 | actin II |
| PBANKA_0936900 | actin-like protein, putative |

**Figure S2. Protein–protein interaction network of Alp5 via the STRING database. (A)** The figure shows protein–protein interactions involving *P. berghei* Alp5a. The network highlights the interactions between Alp5a and other associated proteins. The connectivity and potential functional relationships within the protein interaction network are shown. **(B)** List of proteins that interact with Alp5a. **(C)** Protein association network of *P. berghei* Alp5b. STRING interaction analysis revealed numerous proteins overlapping with the ALP5a interactome. **(D)** List of proteins that interact with Alp5b.

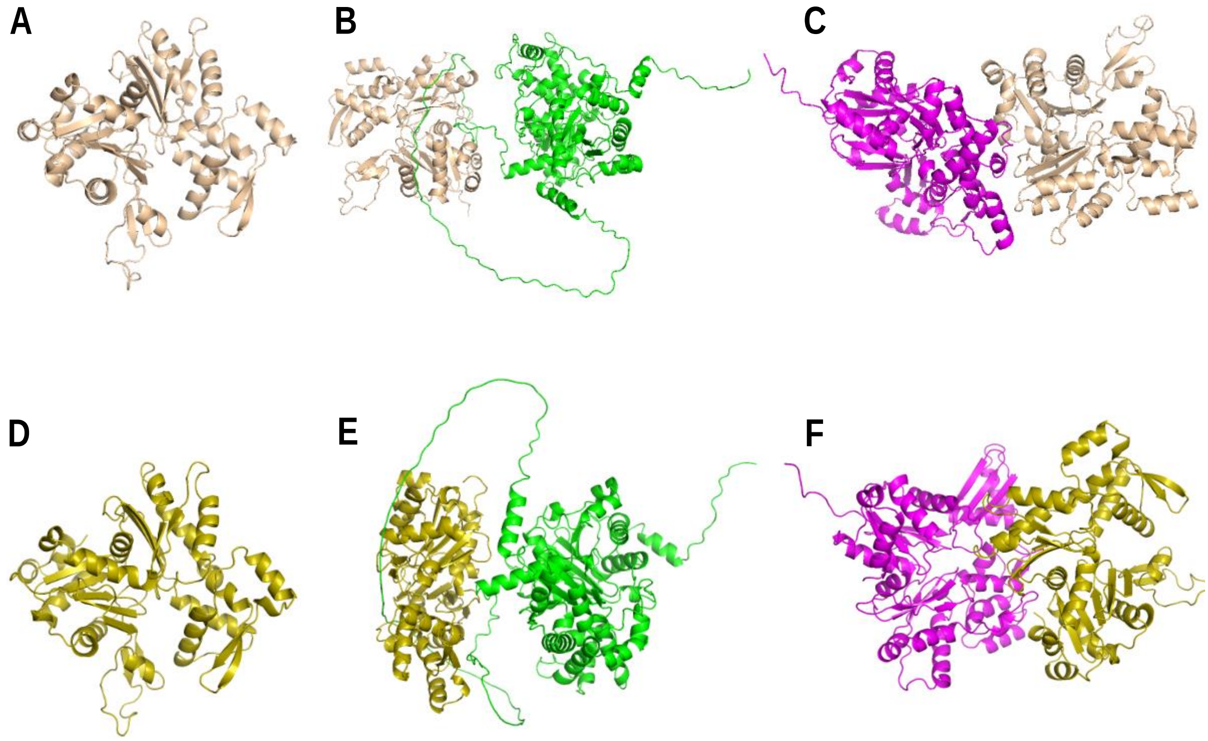


**Figure S3.** Cluspro protein–protein docking studies were performed to predict complex formation, and binding energy calculations were carried out to assess the interaction strength. These results suggest that, compared with Alp5b, Alp5a has a stronger affinity for both actin isoforms. The interaction of Alp5a with actin 1 yields a binding energy of -1187.1 kcal/mol, whereas the interaction with actin 2 shows a favorable binding energy of -1369.1 kcal/mol. In contrast, Alp5b displays weaker binding interactions, with the lowest binding energy of -679.8 kcal/mol for the Alp5b-actin 1 complex and -844.8 kcal/mol for the Alp5b-actin 2 complex. (Alp5a-Green, Alp5b-magenta, Actin 1-wheatish, Actin 2-olive orange). **(A)** Cartoon structure of *P. berghei* Actin 1. **(B)** The figure shows the interaction between *P. berghei* Actin 1 and Alp5a. (**C)** The figure shows the interaction between *Plasmodium berghei* Actin 1 and Alp5b. **(D)** Cartoon structure of *P. berghei* Actin 2. **(E)** Interaction between *P. berghei* Actin 2 and Alp5a. (**F)** The figure shows the interaction between *P. berghei* Actin 2 and Alp5b.

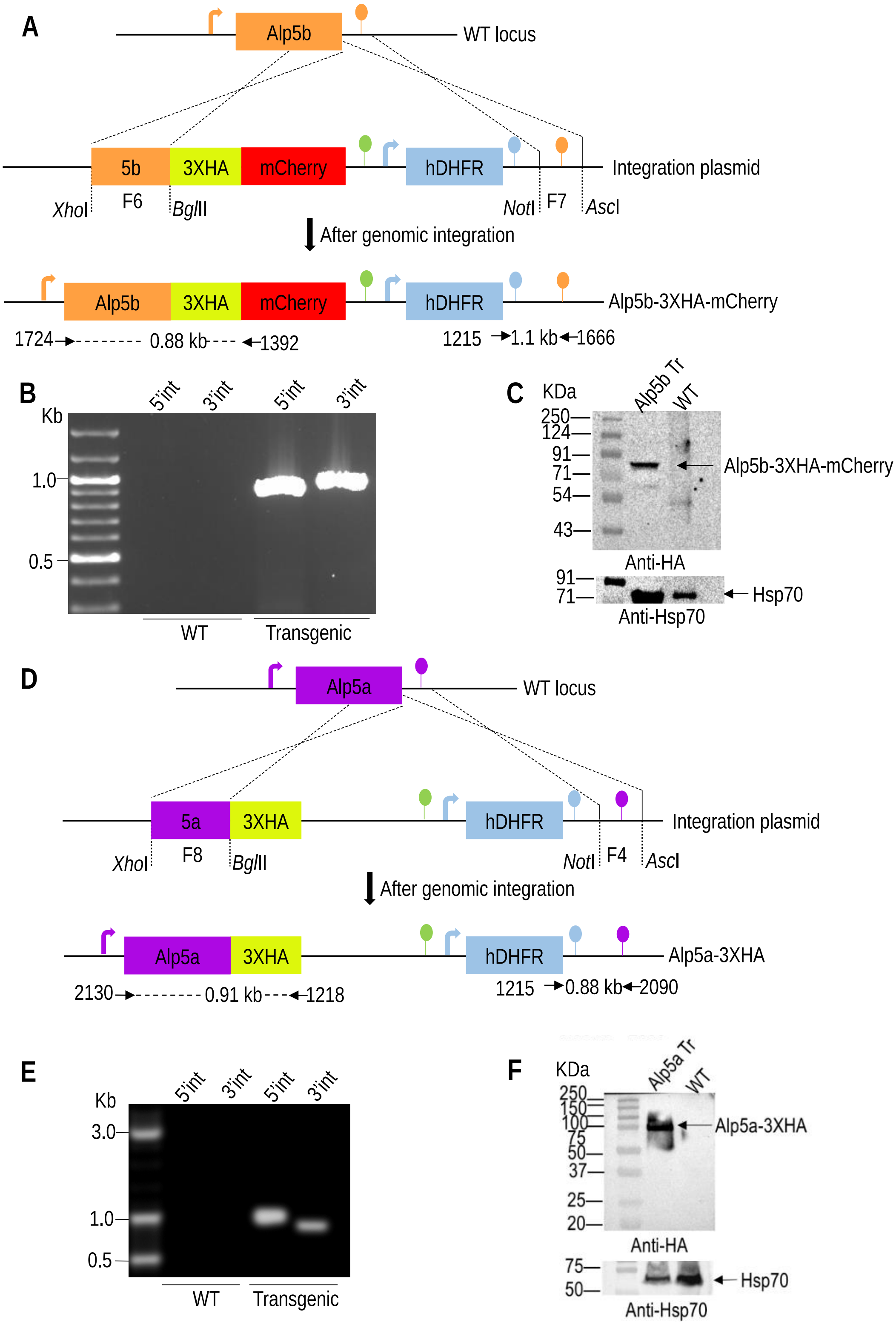


**Figure S4. Generation of transgenic parasites of Alp5b and Alp5a. (A)** To check the expression of Alp5b, the gene was endogenously tagged with 3XHA-mCherry. The arrows and lollipops indicate the 5' and 3' UTRs, respectively. (**B**) Correct site-specific 5' and 3' integration was confirmed via diagnostic PCR via the primer sets 1724/1392 and 1215/1666, respectively. No bands were amplified from WT genomic DNA. (**C**) Western blot analysis of the Alp5b-3XHA-mCherry fusion protein (~77 kDa) revealed via an anti-HA antibody. The blot was reprobed with an anti-Hsp70 antibody as a loading control. **(D)** To check the expression of Alp5a, the gene was endogenously tagged with 3XHA. **(E)** Correct site-specific 5’ and 3’ integration was confirmed via diagnostic PCR via primers 2130/1218 and 1215/2090, respectively. No bands were amplified from WT genomic DNA. (**F**) Western blot analysis of the Alp5a-3XHA fusion protein (70.5 kDa) via an anti-HA antibody. The blot was reprobed with an anti-Hsp70 antibody as a loading control. The expected size bands were observed for the transgenic parasites, whereas no bands were detected in the WT lane.

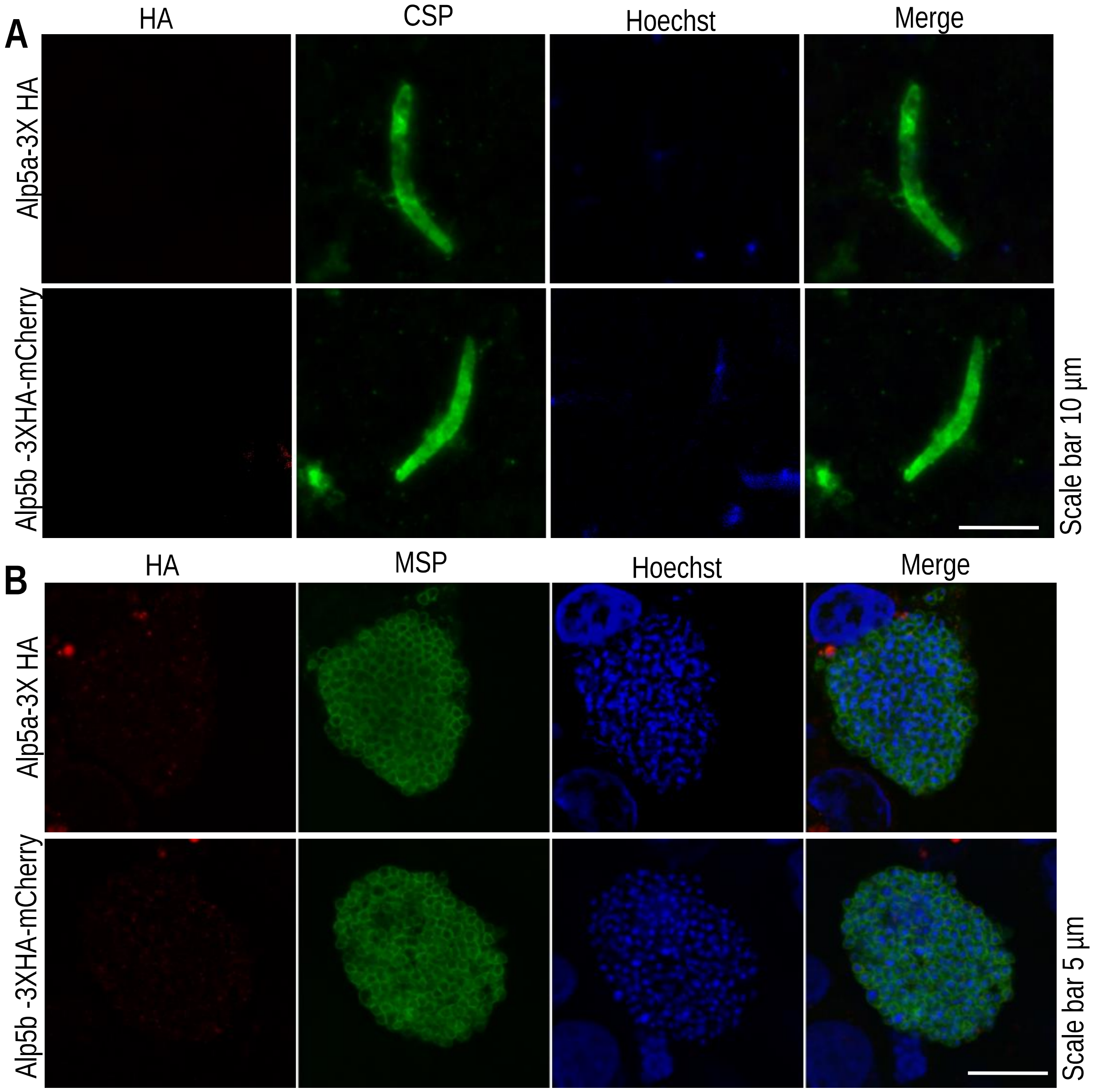


**Figure S5. Alp5a and Alp5b were not expressed in the sporozoite or liver stages. (A)** Expression analysis of Alp5a and Alp5b in sporozoites in tagged parasites. **(B)** Expression analysis of ALP5a and ALP5b in liver stages.

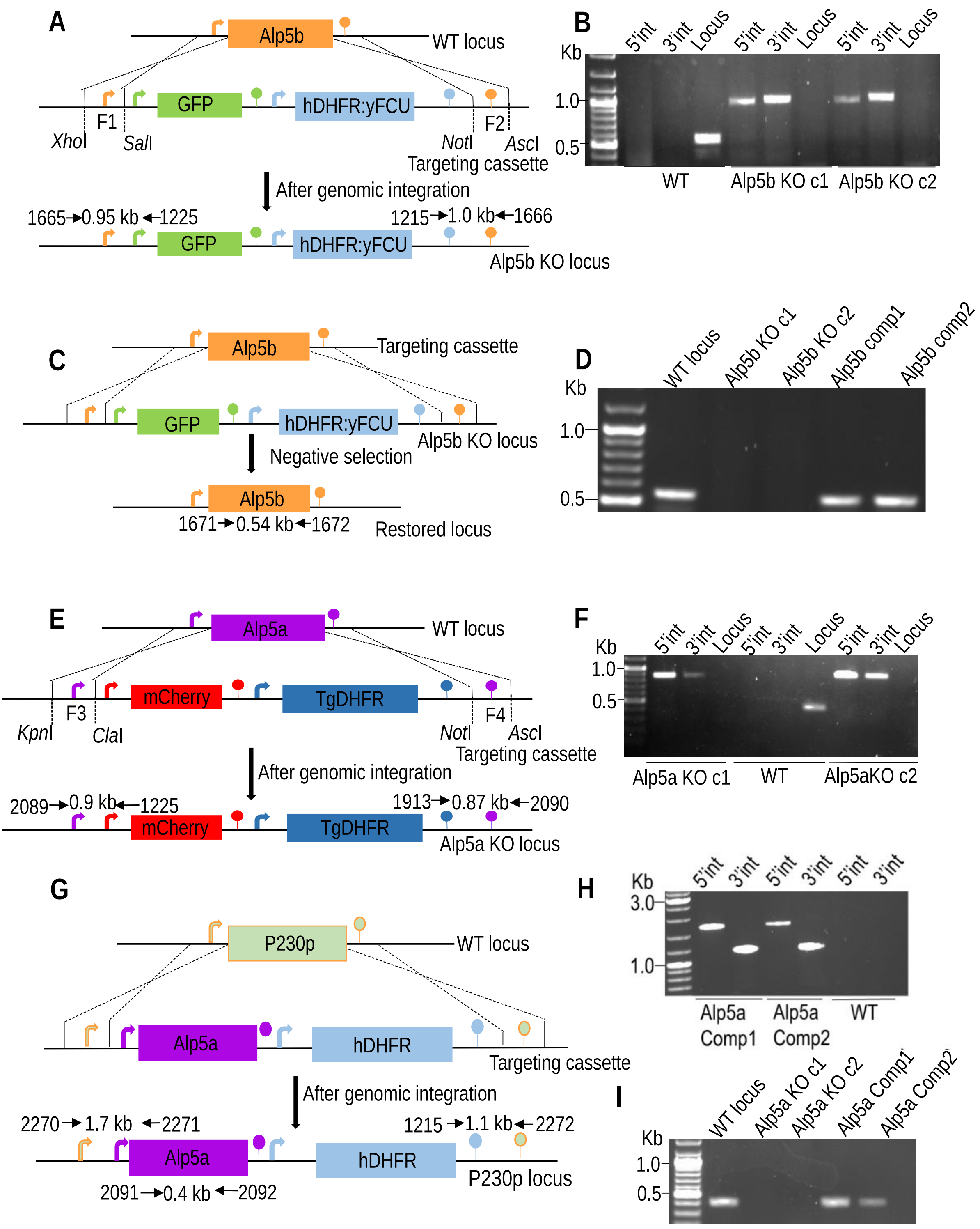


**Figure S6. Generation of Alp5 KO and complemented parasites. (A)** The Alp5b gene was disrupted by double-crossover (DCO) homologous recombination with fragments F1 and F2, as shown in the targeting cassette. The targeting vector pBC-GFP-hDHFR:yFCU consists of the GFP and hDHFR:yFCU cassettes regulated by 5' (arrow) and 3' (lollipop) regulatory sequences. **(B)** Diagnostic PCR showing the correct 5' and 3' integrations via the primer sets 1665/1225 and 1215/1666, respectively. The primers 1671/1672 were used to confirm the absence of the Alp5b locus in the KO parasites, and the band was amplified in the WT parasites but not in the KO parasites. **(C)** Schematic representation of the complementation strategy for reintroducing the

Alp5b gene at the KO locus. After being transfected with the Alp5b-targeting cassette, the parasites were negatively selected with the 5-FC drug. **(D)** Amplification of the locus via the primers 1671/1672 in the WT and complemented lines but not in the KO parasites. **(E)** Similar to Alp5b, Alp5a was disrupted by DCO. The targeting vector pBC-mCherry-tgDHFR consists of the mCherry and tgDHFR cassettes regulated by 5' (arrow) and 3' (lollipop) regulatory sequences. **(F)** Correct site-specific 5' and 3' integrations were confirmed via the primer sets 2089/1225 and 1913/2090, respectively. The absence of a locus in the KO line was confirmed via the primers 2091/2092. **(G)** Schematic showing the Alp5a gene complementation strategy. The Alp5a expression cassette was cloned and inserted into the pBC-P230-hDHFR plasmid, linearized, and transfected into Alp5a KO schizonts, which were subsequently selected via the WR drug. **(H)** Site-specific 5' and 3' integrations of the targeting cassette at the P230 locus were confirmed via diagnostic PCR via primers 2270/2271 and 1215/2272, respectively. **(I)** Amplification of the Alp5a ORF in the complemented line via the primer set 2091/2092.

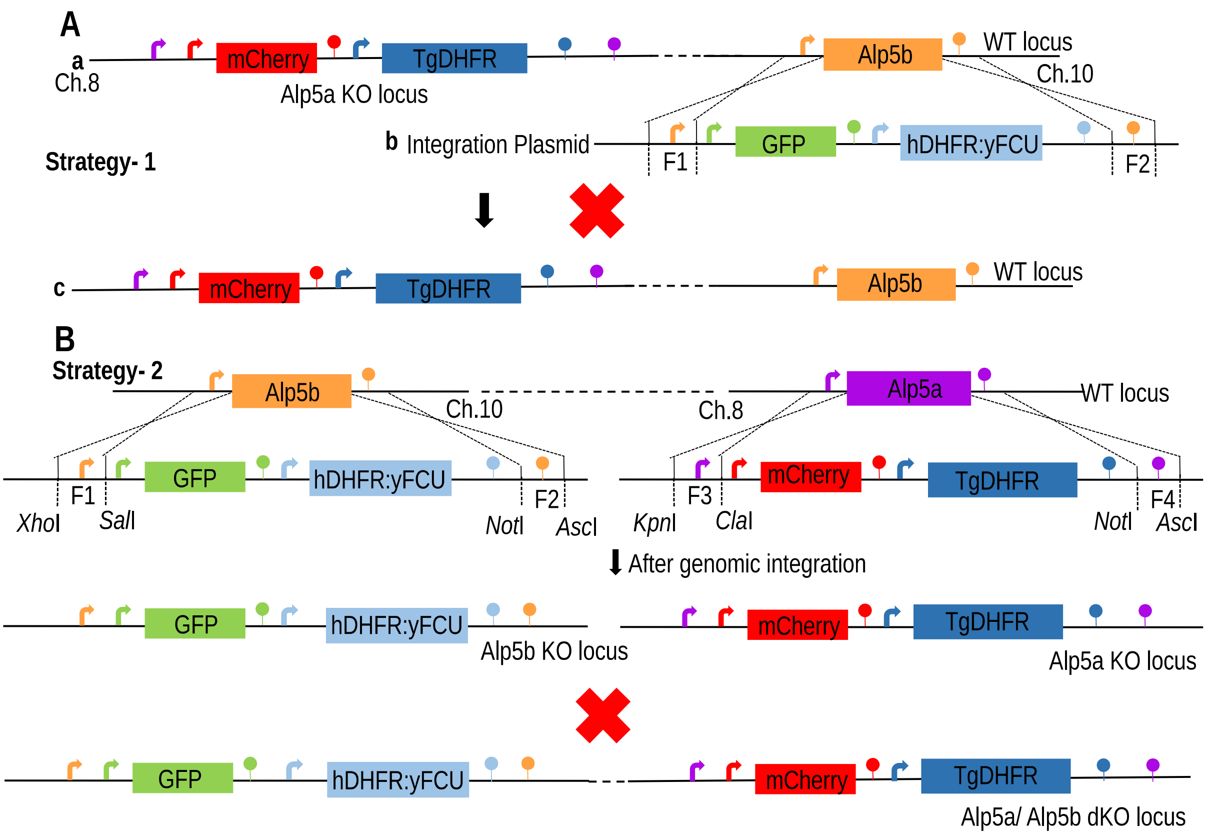


**Figure S7. Attempts to generate Alp5a/Alp5b double-KO parasites. (A)** Schematic representation of the strategy for generating Alp5a/Alp5b double-KO parasites. *Pb*Alp5a KO schizonts were transfected with an Alp5b-targeting cassette. (a) ALP5a locus (b) Recombination at the Alp5b locus. (c) Expected double KO locus. **(B)** Schematic representation of the 2$^{nd}$ strategy. Both targeting constructs were transfected simultaneously into *P. berghei* schizonts.

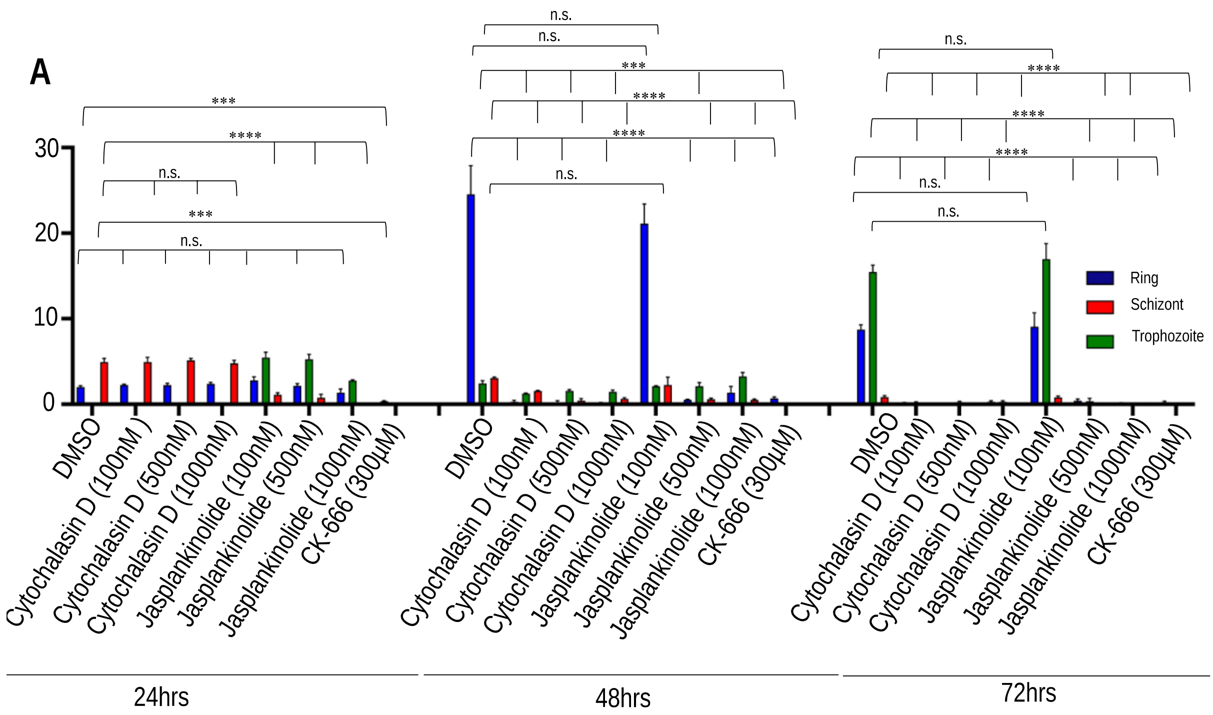


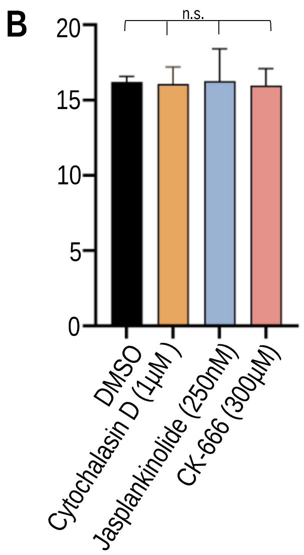


**Figure S8. Effect of inhibitors on the growth of *P. falciparum* and *P. berghei* blood stage parasites. (A)** Counting of ring, trophozoite, and schizont stages in *P. falciparum* cultures after treatment. There was no significant difference in the number of schizonts between the cytochalasin D-treated group and the DMSO-treated group at 24 h (P= 0.9488). The number of schizonts significantly decreased in the jasplankinolide (****P<0.0001)- and CK-666 (***P= 0.003)-treated groups. Cytochalasin D and jasplankinolide did not affect the ring stage at 24 h (P=0.0638). The number of ring stages significantly decreased in the CK-666-treated groups (***P=0.0008). There were fewer schizont numbers at 48 h in all the treated groups (****P<0.0001). The number of ring stages observed at 48 h also decreased significantly (****P<0.0001). The number of trophozoites was significantly lower in all the treated groups (***P=0.0001). All the stages observed at 72 h were significantly reduced in all the treated groups (****P<0.0001). **(B)** *P. berghei* blood cultures were treated with the inhibitors cytochalasin D (1 µM), jasplankinolide (250 nM) and CK-666 (300 µM). Parasitemia was comparable between the treated and control groups (P= 0.9984; Brown-Forsythe ANOVA).

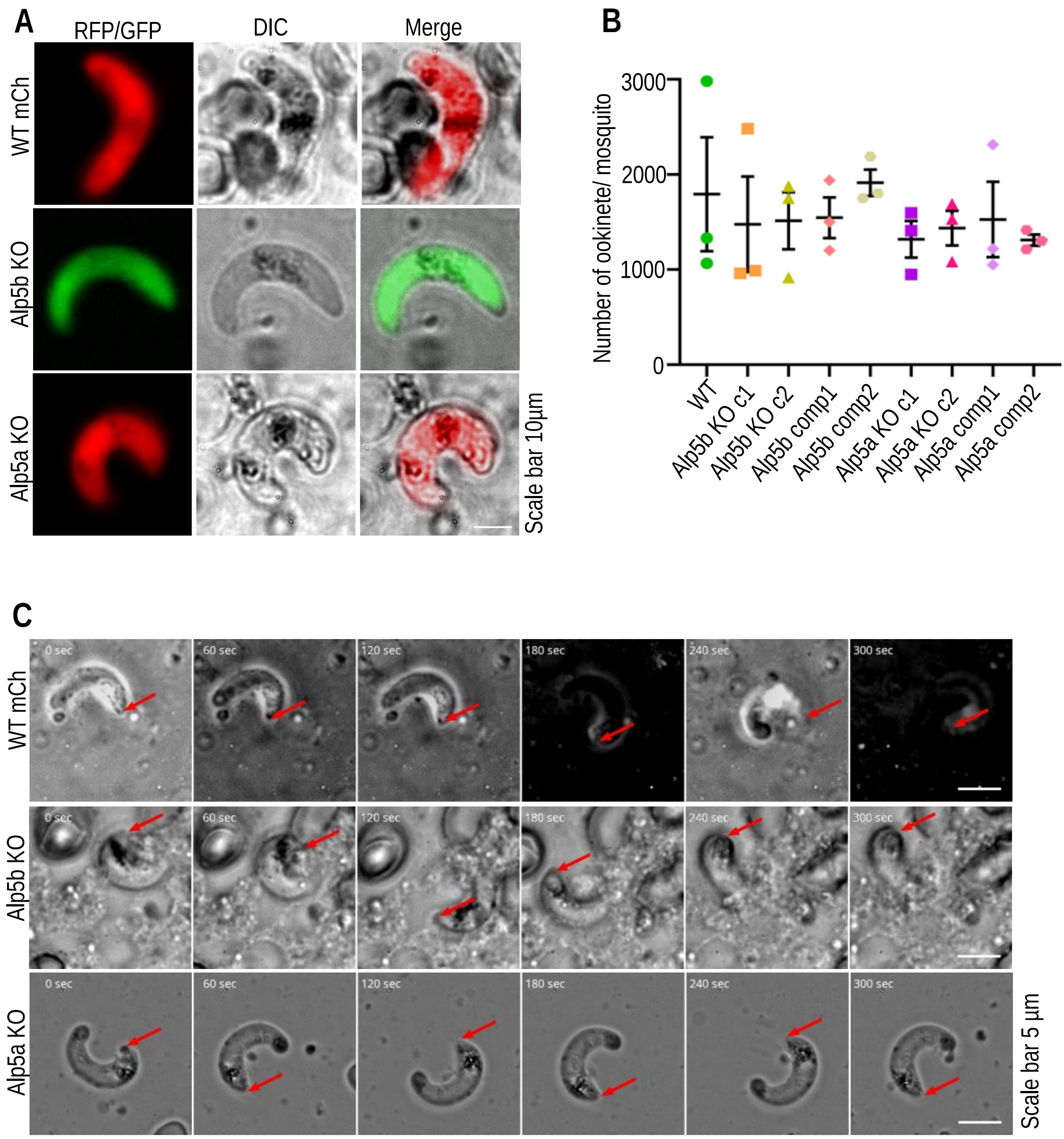


**Figure S9. Hypoploidy of male gametes does not affect the viability of ookinetes (A)** The morphology of Alp5 KO ookinetes was similar to that of WT parasites. **(B)** There was no difference in the number of ookinetes between WT and Alp5 KO parasites (P= 0.9275, one-way ANOVA). Ookinete numbers from 109 (WT-mCh), 72 (Alp5b cl1), 75 (Alp5b cl2), 110 (Alp5b comp1), 65 (Alp5b cl2), 75 (Alp5a cl1), 105 (Alp5a cl2), 65 (Alp5a comp1), and 85 (Alp5a comp2) midguts were observed. **(C)** Ookinetes were allowed to glide for 5 minutes with 61 loops using Matrigel. No difference in gliding was observed between WT and Alp5 KO parasites.

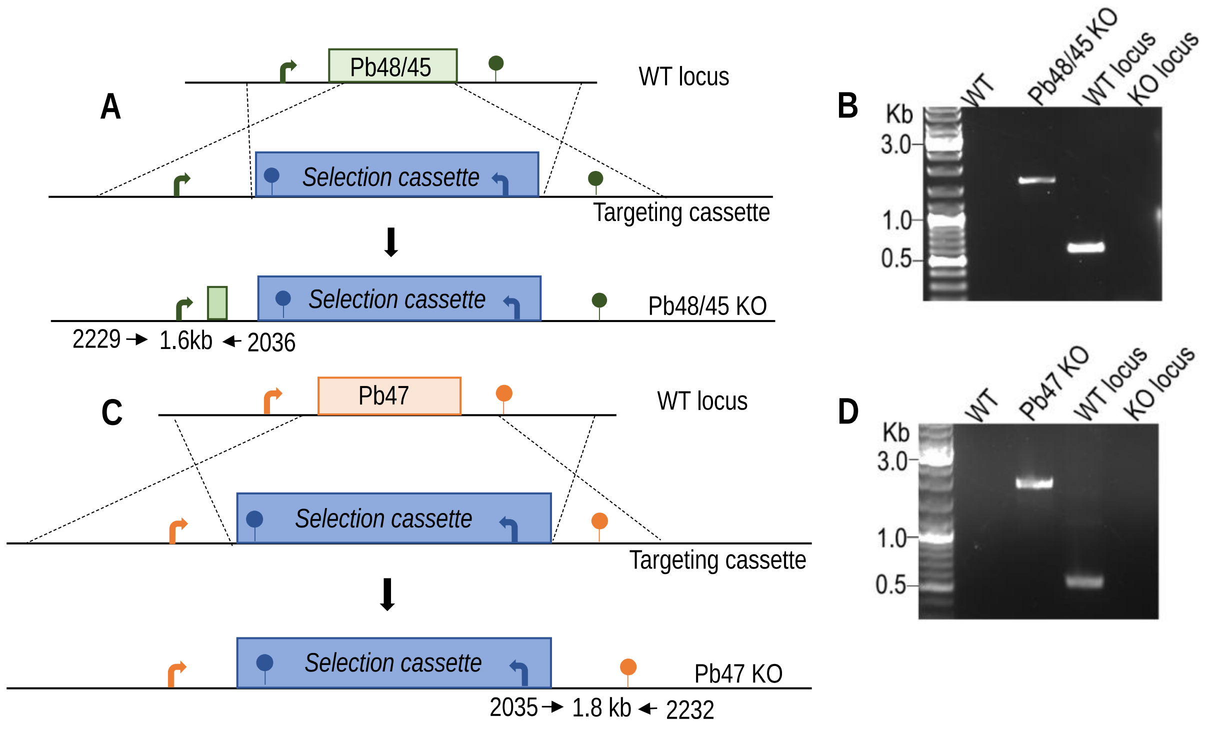


**Figure S10. Generation of male and female gamete defective parasite lines. (A)** The strategy used to generate a male gamete-defective Pb48/45 KO parasite line. **(B)** Genotyping confirmed the correct site-specific integration using primers 2229/2036. The ORF was amplified from WT genomic DNA but not from Pb48/45 KO parasites via primers 2230/2231. **(C)** Strategy used to generate a female gamete-defective Pb47 KO parasite line. **(D)** Site-specific integration PCR using primers 2232/2035. The ORF was amplified from WT genomic DNA but not from Pb47 KO parasites via primers 2233/2234.

| S. No. | Cross between | Possible combinations of gamete formed | Phenotype of oocyst in midguts | Inference |
|---|---|---|---|---|
| 1. | WT mCh × ALP5b KO | ♂ ♀<br>♀ ♂<br>♂ ♀<br>♂ ♀ | 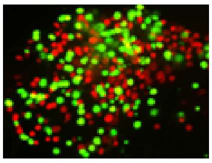 | Phenotype restored |
| 2. | WT GFP × ALP5a KO | ♂ ♀<br>♀ ♂<br>♂ ♀<br>♀ ♂ | 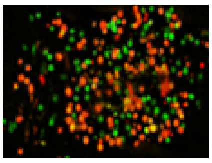 | Phenotype restored |
| 3. | Pb47 KO × ALP5b KO | ♂ ♀<br>♀ ♂<br>♂ ♀<br>♂ ♀ | 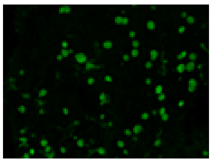 | Phenotype restored |
| 4. | Pb48/45 KO × ALP5b KO | ♂ ♀<br>♀ ♂<br>♂ ♀<br>♂ ♀ | 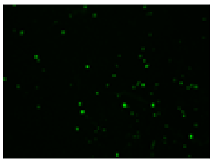 | No change |
| 5. | Pb47 KO × ALP5a KO | ♂ ♀<br>♀ ♂<br>♂ ♀<br>♂ ♀ | 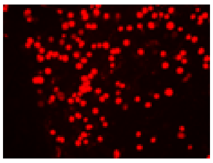 | Phenotype restored |
| 6. | Pb48/45 KO × ALP5a KO | ♂ ♀<br>♀ ♂<br>♂ ♀<br>♂ ♀ | 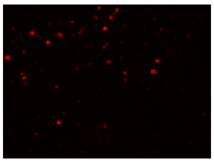 | No change |

**Figure S11. Summary of the genetic cross experiments.** Genetic crosses with WT or female gamete-defective parasite lines restored the KO phenotype. There was no change when the KO was crossed with the male gamete-defective parasite line.

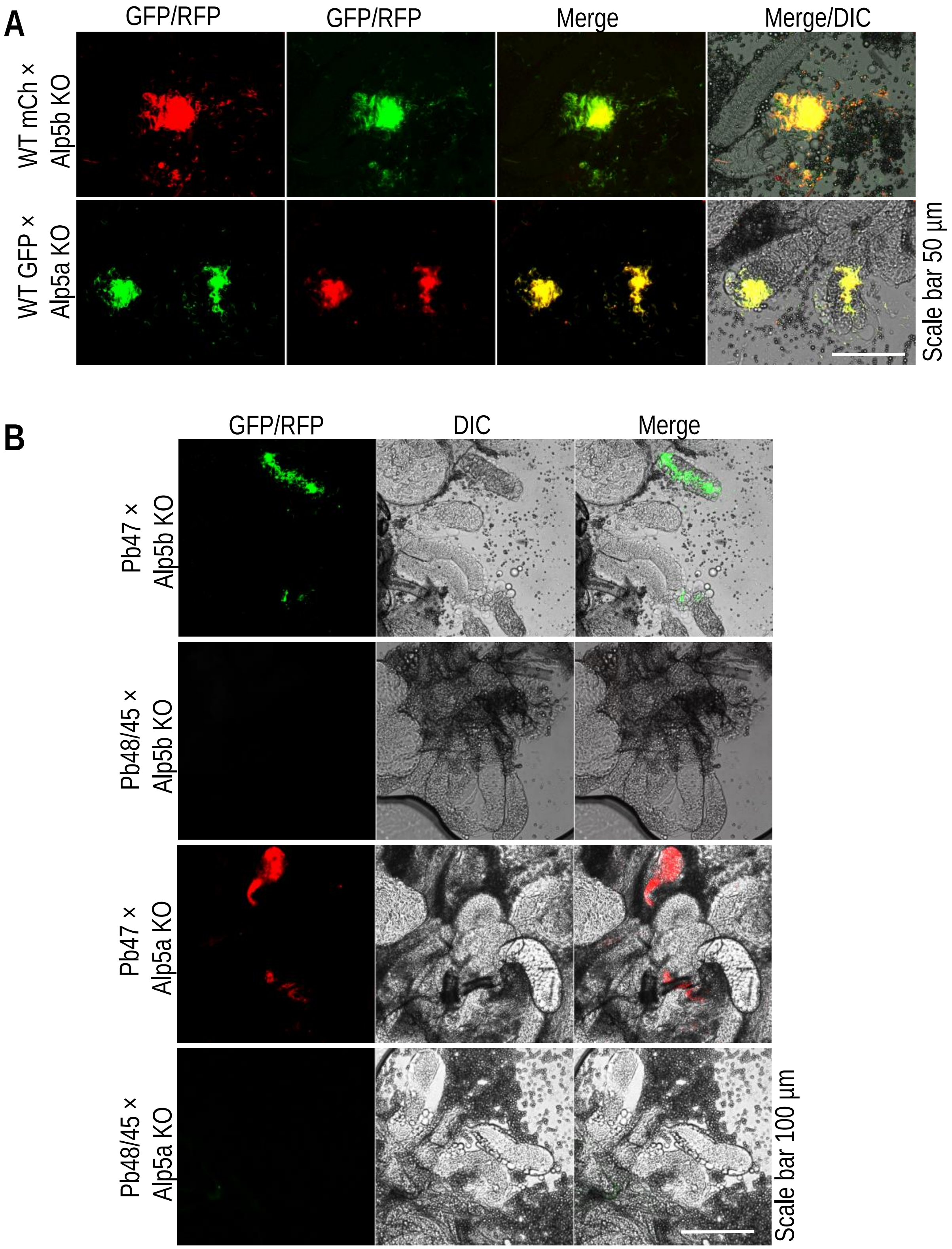


**Figure S12. Infected salivary glands showing the sporozoite load. (A)** Observation of the salivary gland sporozoite load after genetic crosses with WT-mCh×Alp5b KO and WT-GFP×Alp5a KO. **(B)** The salivary gland sporozoite load in Alp5 KO lines after genetic crosses with male (Pb48/45 KO) or female (Pb47 KO) gamete-defective lines.

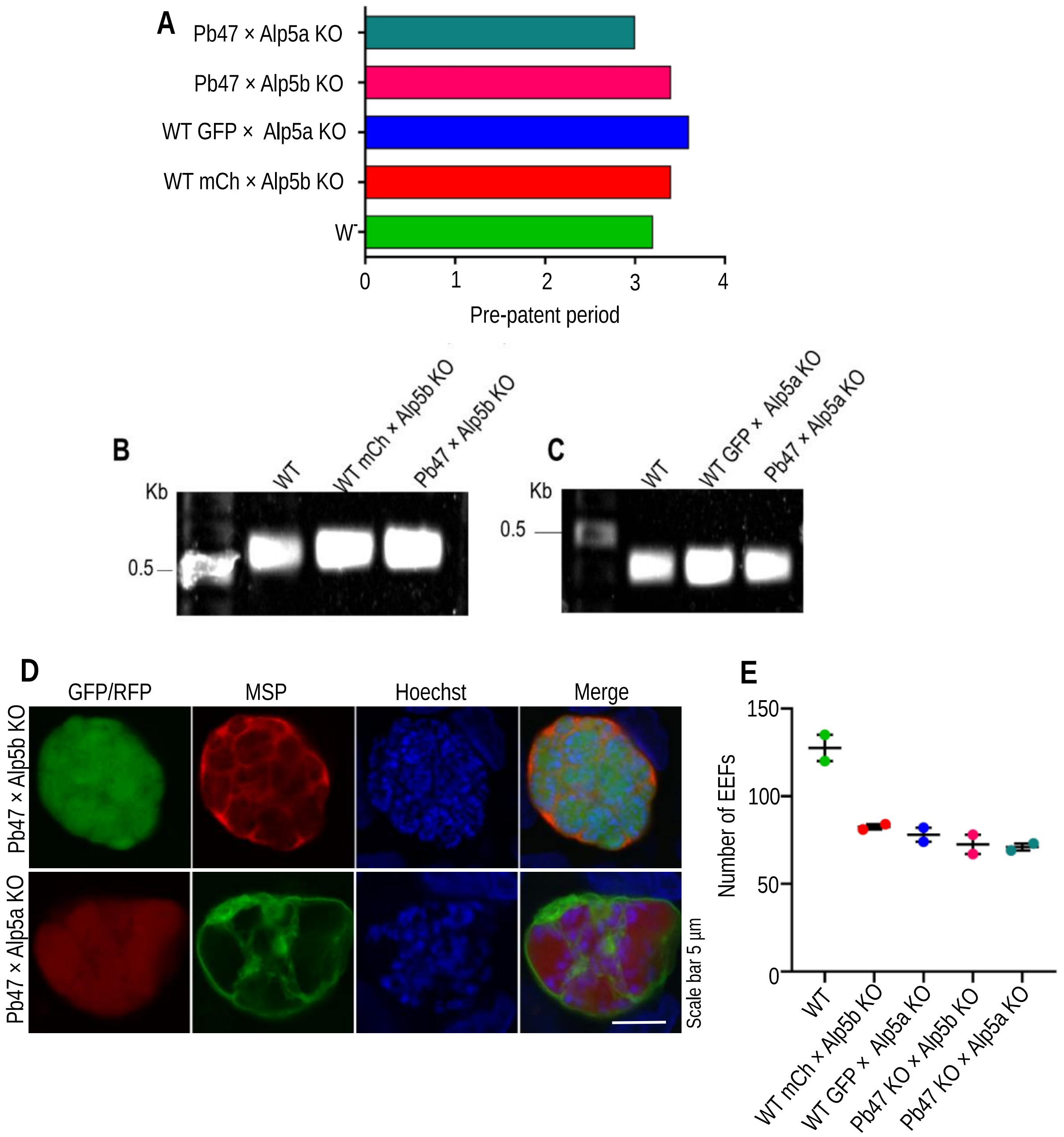


**Figure S13. In vivo and in vitro infectivity of sporozoites. (A)** Salivary gland sporozoites were injected into C57BL/6 mice, and the pre-patent period was observed. Genetic crosses with WT or female gamete-defective Pb47 KO lines restored the infection of Alp5 KO parasites **(B and C).** The Alp5 ORF was amplified in the KO lines after genetic crosses with WT or gamete-defective lines. **(D)** Sporozoites invade hepatocytes and develop into EEFs. **(E)** EEF counts at 65 h. Only GFP- or mCherry-expressing EEFs were counted.

**Table S1.** List of primers used in this study.

| Primer | Primer sequences | Restriction site |
|---|---|---|
| 1611 | GTCGAC TCCTATTCTTATCAAATTTACCCTA | *Sal*I |
| 1612 | GTCGACACGGGAACATTTTTCCTGTCC | *Sal*I |
| 1613 | GCGGCCGCTGGGCAAACATCCTATAAAGG | *Not*I |
| 1614 | GGCGCGCCGTGCTAGAACAATTTGGATATAT | *Asc*I |
| 1665 | GATTATGCGGGTATTCATATTTGCATC | NA |
| 1666 | TAACTATGAAAGAAGGTGAAAATGAAAATAGTG | NA |
| 1671 | GAATCAAATAGTTATGTAAGTCTA | NA |
| 1672 | TAGGATGTTTGCCCAAATAAAAA | NA |
| 1675 | CTCGAGGAATCAAATAGTTATGTAAGTCTA | *Xho*I |
| 1676 | AGATCTTAGGATGTTTGCCCAAATAAAAA | *Bgl*II |
| 1677 | GCGGCCGCAGGTGTAGAGAACAAGAAAATA | *Not*I |
| 1724 | TTCCTGTGGCTGAAAATAATG | NA |
| 1895 | TATAGGGCGAATTGGGTACCATAATTCGATTGTGTATTTTTATT | *Kpn*I |
| 1896 | GTATATTTTCCATCGATGCTCATTTTATTTTACATTTATTA | *Cla*I |
| 1897 | TGCAAGCTTGCGGCCGCTAATTTTTTTTTTGCATATAAGAAG | *Not*I |
| 1898 | ATTACGCCAGGCGCGCCACTTTCATCTTATTATATTTGTTTC | *Asc*I |
| 2089 | GTCTATGCTTTTCTATCATGC | NA |
| 2090 | AGCAGGGGATGGTTATTTTA | NA |
| 2091 | CGGACTCATATAATAATTTGC | NA |
| 2092 | ATACAATAACTTTCCTTGTAAG | NA |
| 2128 | CGGGCCCCCCCTCGAGCTCCCCAAAATAAAGATGAATCTG | *Xho*I |
| 2129 | CGTATGGGTAAGATCTATACAATAACTTTCCTTGTAAGCTC | *Bgl*II |
| 2130 | CCGACTTAACTAGCATATATT | NA |
| 2268 | CAGTCGACGGTATCGATATAATTCGATTGTGTATTTTT | *Cla*I |
| 2269 | TGGGCTGCAGGAATTCATATAAATATATCACATACTTTCA | *EcoR*I |
| 2270 | GCTTGACCATGATTACGCC | NA |
| 2271 | TATTATAATTAATGTAAGCG | NA |
| 2272 | ATCAAGATAAATATACCATAAGT | NA |
| 2229 | TAATTTAGGTATGAAATGTAGG | NA |
| 2230 | GATGGAAGAAGTGCATTAGT | NA |
| 2231 | CTGGAATTATAGTACCAGGG | NA |
| 2232 | GTTCCCAAAAAAGTAGAGCA | NA |
| 2233 | CTGAAGACAGCGCACACA | NA |
| 2234 | GCCACCAACATTGCTAAAAA | NA |
| 2035 | CATACTAGCCATTTTATGTG | NA |
| 2036 | CTTTGGTGACAGATACTAC | NA |
| 1215 | GTTGTCTCTTCAATGATTCATAAATAG | NA |
| 1225 | TTCCGCAATTTGTTGTACATA | NA |
| 1392 | GCCCTCCATGTGCACCT | NA |
| 1218 | TACAACAAAAGGAGGTACAC | NA |
| 1913 | ACCAACTCAATTTAATAGATGT | NA |
